# A Model-Based Hierarchical Bayesian Approach to Sholl Analysis

**DOI:** 10.1101/2023.01.23.525256

**Authors:** Erik Vonkaenel, Alexis Feidler, Rebecca Lowery, Katherine Andersh, Tanzy Love, Ania Majewska, Matthew N Mccall

## Abstract

Due to the link between microglial morphology and function, morphological changes in microglia are frequently used to identify pathological immune responses in the central nervous system. In the absence of pathology, microglia are responsible for maintaining homeostasis, and their morphology can be indicative of how the healthy brain behaves in the presence of external stimuli and genetic differences. Despite recent interest in high throughput methods for morphological analysis, Sholl analysis is still the gold standard for quantifying microglia morphology via imaging data. Often, the raw data are naturally hierarchical, minimally including many cells per image and many images per animal. However, existing methods for performing downstream inference on Sholl data rely on truncating this hierarchy so rudimentary statistical testing procedures can be used. To fill this longstanding gap, we introduce a fully parametric model-based approach for analyzing Sholl data. We generalize our model to a hierarchical Bayesian framework so that inference can be performed without aggressive reduction of otherwise very rich data. We apply our model to three real data examples and perform simulation studies comparing the proposed method with a popular alternative.

## 1. Introduction

It has been shown that microglia are key players in countless brain pathologies including neurodegenerative disorders, traumatic brain injury, and psychiatric diseases (Sierra and *others*, 2019; Prinz *and others*, 2019; Gomez-Nicola and Perry, 2014). As the main immune cells in the central nervous system, microglia respond to these pathologies in a myriad of ways. Alongside reactive behavior, microglia may also have a direct impact at the onset of several diseases. For example, recent genome-wide association studies showed that genes which are risk factors for Alzheimer’s disease are largely expressed in microglia rather than in other brain cell types (Hemonnot *and others*, 2019). Other specific pathologies which involve the microglia include glioma, strokes, multiple sclerosis, Parkinson’s disease, autism, and schizophrenia (Hambardzumyan *and others*, 2016; Patel *and others*, 2013; Long-Smith *and others*, 2009; Takano, 2015; Monji *and others*, 2009; Bogie *and others*, 2014). Further, studies have also linked microglia function to various lifestyle factors such as stress, diet, and alcohol consumption (Tynan *and others*, 2010; Johnson, 2014; Marshall *and others*, 2013).

A primary reactive behavior of microglia is to change their morphological phenotype. Homeostatic microglia are ramified cells, characterized by a number of highly branched processes extending from a central soma. In response to the presence of either pathological or physiological stimuli, microglia can re-organize these processes to change their number, shape, and distribution, resulting in a broad spectrum of morphological phenotypes. Theses morphological changes are a dynamic process which potentially differ depending on the stimulus, environmental context, and the stage of the microglial response (Franco and Fernández-Suárez, 2015; Tang and Le, 2015). While it is challenging to make direct inferences about microglial function based purely on morphological changes (Paolicelli *and others*, 2022), morphology remains an important indicator of changes in microglial function in many different physiological and pathological settings.

Though there has been interest in high-throughput methods for analyzing microglial morphology (Colombo *and others*, 2022; Heindl *and others*, 2018), often studies rely on simple analysis methods implemented in freely available software, such as Sholl analysis (Sholl, 1953), as a means to quantify cell morphology. Despite the popularity of Sholl analysis, methods for performing inference on cell morphology using Sholl data are extremely limited. Though Sholl analysis is able to capture a wide range of morphological changes, current methods struggle to take advantage of all available information. We aim to fill this gap by proposing novel methods for performing inference using Sholl data.

In this article, we propose a fundamentally different inference procedure for Sholl data. Specifically, we propose a model based approach using biologically meaningful parameters. We adopt a hierarchical Bayesian framework, which can easily capture variation at each level of the experimental hierarchy. Further, the model is parameterized so that Sholl curve summaries commonly used in the literature can be retrieved directly from the model parameters. This allows investigators to perform inference using all of the data available to them and incorporate variation at each level of the experimental hierarchy.

## 2. Methods

### 2.1 Existing Methods

To perform Sholl analysis, one constructs concentric circles around the soma of a cell, the smallest containing the soma, and the largest containing the entire process arbor, i.e. the entire cell. Then, the number of times any process crosses each circle is counted. A Sholl curve (Figure S1 A) is constructed by plotting the counts for each circle against the corresponding radii.

Currently, some of the most popular morphological analysis methods are based on the analysis of Sholl curves. Most existing methods involve aggressive data reduction or transformation so that basic statistical procedures can be used. There are two primary avenues for analyzing these data: transformation-based and summary-based methods.

Transformation-based methods involve linearizing Sholl curves so that ordinary least squares can be applied, the most common being the *semi-log* and *log-log* methods. For the *i^th^* concentric circle, let *x_i_* be the radius, *A_i_* be the area, and *y_i_* be the number of intersections. Then the semi-log regression model is given by 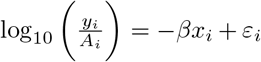, where *ε_i_* ~ *N*(0,1). Similarly, the log-log model is given by 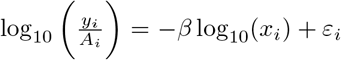, where *ε_i_* ~ N(0,1). The parameter *β* is called Sholl’s regression coefficient, which is often interpreted as the decay rate of the number of branches with distance from the soma (Sholl, 1953; Milosević and Ristanović, 2006).

These linearizations can be quite poor, typically resulting in the transformed Sholl curve oscillating about the fitted linear curve. Even when the linearization appears reasonable, the typical oscillation is still apparent (Figure S1 B). Additionally, we commonly have access to many cell images, often in some nested hierarchical structure induced by experimental design, so a single linearization technique may not be appropriate for all available data. Even if a model is reasonably chosen, we are still limiting our analysis of very rich data to a single parameter model.

Another strategy involves reducing the Sholl curve into a summary statistic, which can be passed to a hypothesis testing procedure. Some previously proposed Sholl curve summaries are:

- Branch Maximum: the maximum number of crossings across all radii
- Critical Value: the radius at which the maximum number of crossings is observed
- Schoenen Ramification Index: the branch maximum divided by the number of branches originating at the soma
- Area Under the Curve
- Full Width Half Max: the width of the curve at half the maximum number of crossings

Often, there is some amount of averaging that occurs before these summaries are calculated.

Hierarchical data are collapsed at the subject-level so that each subject only has a single, aggregated Sholl curve. This aggregate curve is typically the point-wise mean of each cell-level curve associated with that subject. Inference is commonly performed on Sholl curve summaries using ANOVA so that group differences and interactions can be tested.

### 2.2 The Sholl Curve Model

We start by specifying our model for a single Sholl curve. Let 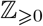 and 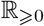 denote the set of integers and real numbers greater than or equal to 0, respectively. Then a Sholl curve is the pair (*X*, *Y*), where 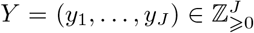 are the process crossings corresponding to the concentric shells of radius 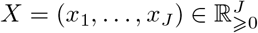, where *x_1_* < … < *x_J_*. The model is then given by:

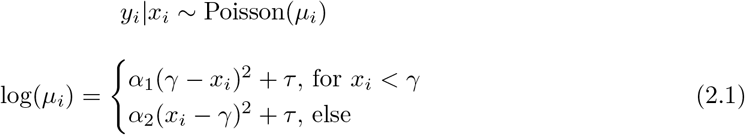

where *α*_1_, *α*_2_ < 0, 0 < γ < *x_J_*, and *τ* > 0. This is essentially a change-point generalized non-linear model assuming a Poissonian random component with canonical link function. Intuitively, the log transform of Sholl curves are approximately asymmetric quadratics (Figure S2), which we directly model in the log-mean function. Since Sholl curves are count data, a Poissonian random component with canonical log link is a natural approach.

We think of this model as a combination of a “growth-curve” and a “decay-curve”, which are separated by the change-point *γ*. The parameters *α*_1_ and *α*_2_ control the growth and decay curves, respectively. The maximum of the fitted curve is given by (*γ*, *e^τ^*), allowing us to directly estimate the critical value and branch maximum. We can also retrieve the *y*-intercept of the estimated curve as *α*_1_ *γ*^2^ + *τ*, which is interpreted as the expected number of processes originating from the soma. Changes in the mean model as each parameter varies can be seen in Figure 1.

**Fig. 1.**
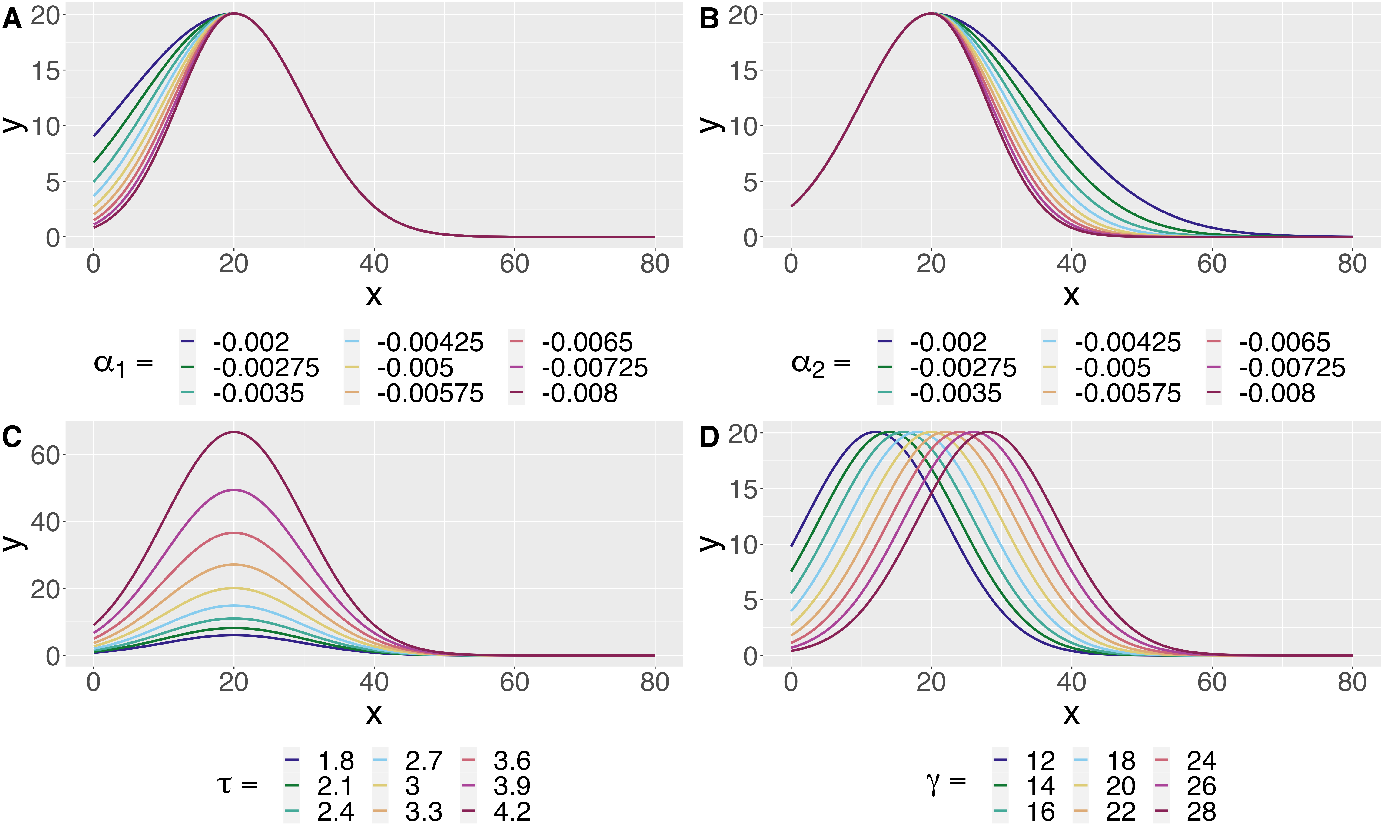
The mean model induced by Equation 2.1 as each parameter varies. **A**: The growth parameter *α*_1_ controls the behavior of the curve before the change-point. **B**: The decay parameter *α*_2_ controls the behavior of the curve after the change-point. **D**: The parameter *τ* controls the branch maximum of the fitted curve via *e^τ^*. **C**: The parameter *γ* controls the critical value, i.e. the change-point, of the fitted curve.

Microglia experiments often contain many images per animal and many cells per image, so a natural extension of the single curve model is a Bayesian hierarchical framework. We demonstrate this extension in three hierarchical models which are later applied to real data examples. Though specifically tailored to our examples, the three models we showcase can be used in many practical settings with minor adjustments.

### 2.3 Model 1

The first model extends the single curve model to a nested hierarchical structure with four levels. In the applied example, the four levels correspond to population, animal, image, and cell, which are nested in that order. We assume parameters for each level are independently sampled from the corresponding distribution in the next highest level. For instance, the model parameters in Equation 2.1 at the cell level are independently sampled from image-level distributions.

The hierarchical structure of this model is displayed in Figure 2. The population-level parameters seen in Figure 2 are defined as *μ* = (*μ*_*α*_1__, *μ*_*α*_2__, *μ_γ_*, *μ_τ_*). Then for the *l^th^* animal, we define *ω_l_* = (*ω*_*α*_1_*l*_ *ι*_*α*_2_*l*_, *ω_γl_*, *ω_τ_*, *l*) and 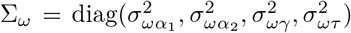. Parameters corresponding to the *k^th^* image of animal *l* are defined as *ψ_kl_* = (*ψ*_*α*_1_*kl*_, *ψ*_*α*_2_*kl*_, *ψ_γkl_*, *ψ_τkl_*) and 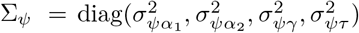. In Figure 2, cell-level parameters for image *k* of animal *l* are vectorized as Θ_*kl*_ = (*θ*_1*kl*_,…, *θ_J_kl_kl_*), so that parameters for cell *j* in image *k* of animal *l* are given by *θ_jkl_* = (*θ*_*α*_1_*jkl*_,*θ*_*α*_2_*jkl*_,*θ_γjkl_*,*θ_τjkl_*) and 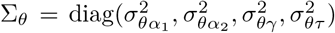. We also vectorize the Sholl curve process crossings for cells in image *k* of animal *l* as *Y_kl_* = (y_1*kl*_,…, y_*J_kl_kl*_), where y_*jkl*_ = (*y*_1*jkl*_,…, *yN_jkl_jkl*) denote process crossings for cell *j* in image *k* of animal *l*.

**Fig. 2.**
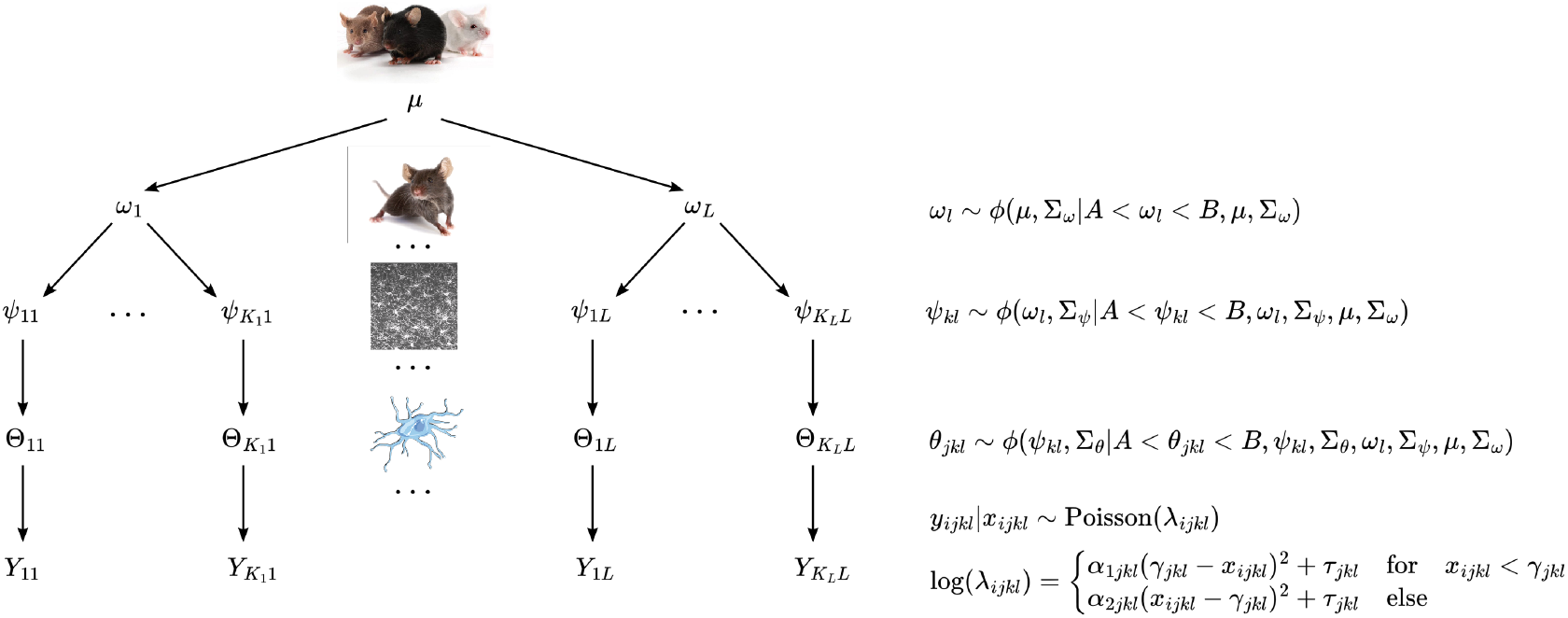
Hierarchical structure for model 1. We assume parameters at any level are randomly sampled from the corresponding distribution in the next highest level. Here, *ϕ*(*) denotes the Gaussian distribution, *μ* denotes population-level parameters, *ω* denotes animal-level parameters, *ψ* denotes image-level parameters, *θ* denotes cell-level parameters, and *Y* denotes Sholl curve process crossings. For a given parameter *, Σ_*_ denotes variance parameters for the corresponding Gaussian. Gaussian priors are truncated via *A* and *B* to enforce the parameter constraints of Equation 2.1.

We truncate normal distributions *ϕ* at each level of the hierarchy according to the parameter space of Equation 2.1. The lower bound of the parameter space is *A* = (−∞, −∞, 0, 0) and the upper bound is 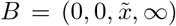, where 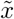 is the least upper bound on the support of the Sholl curves. As suggested in Gelman (2006), we assume a half-t prior on all standard deviation parameters.

### 2.4 Model 2

The second model is a truncation of the first, which we apply to Sholl data that has been aggregated at the animal-level. This model contains population, group, and animal levels, nested in that order. Notably, we model group and interaction effects for each parameter in Equation 2.1 so that group-level inference can be performed. This model allows for two categorical variables, each with two groups.

We can see the hierarchical structure of the model in Figure 3. We include effects at the group level via additive terms on the group level mean parameter. Denoting groups for the first categorical variable as either ND or MD,

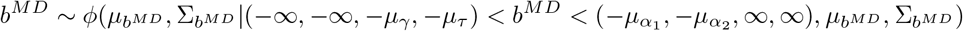

is interpreted as a shift in the group mean for group MD. Similarly denoting groups for the second categorical variable as either I or C,

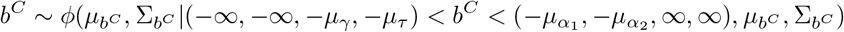

is interpreted as a shift in the group mean for group C. The interaction effect is given by

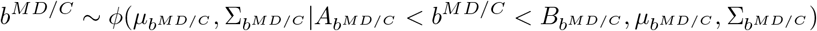

where

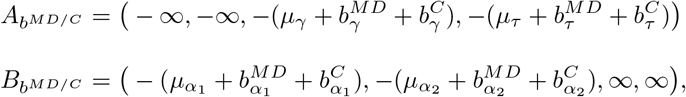

which is interpreted as a shift in the group mean for group MD/C.

**Fig. 3.**
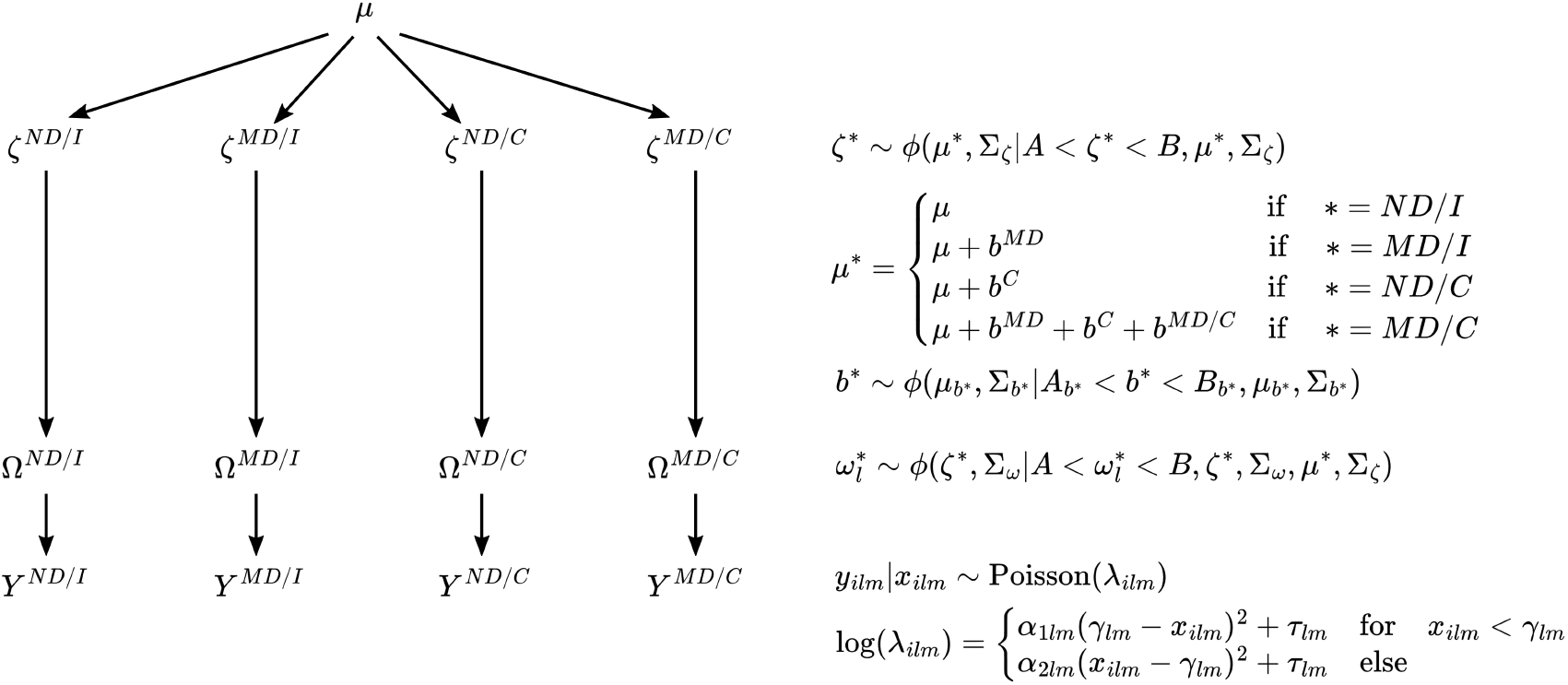
Hierarchical structure for Model 2. Denote groups in the first categorical variable as either ND or MD, and the second as either I or C. Notation is consistent with Figure 2 except, for some combination of groups *, *ζ*^*^ denotes group-level parameters, 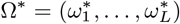 denotes animal-level parameters, and *Y*\* = (*y*_1*lm*_,…, *y_Nlm_*) denotes Sholl curve process crossings. Additionally, group combination is indexed by *m* and we model group level effects as additive terms *b*\* on the mean parameter for group level distributions.

The bounds on these effects are set to constrain the group level means within (*A*, *B*), i.e. the support of the truncated normal distributions. We assume truncated normal hyper-priors on mean parameters for each effect, where the truncation is identical to the corresponding effect bounds. As before, we assume half-t priors on all standard deviation parameters in the model, including half-t hyper-priors for the effect standard deviations.

### 2.5 Model 3

The third model generalizes the second by incorporating cell-level data. Specifically, this model contains population, genotype, animal, and cell levels in that order. We also show how effects can be modeled at multiple levels of the hierarchy by including a genotype-level effect, and cell-specific effects shared across animals.

The full model is shown in Figure 4. At the genotype level, we add an effect for knockout (KO) as an additive term on the mean parameter for *ξ^KO^*. Specifically,

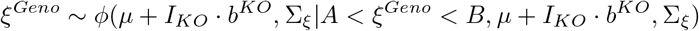

where

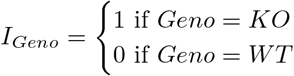

**Fig. 4.**
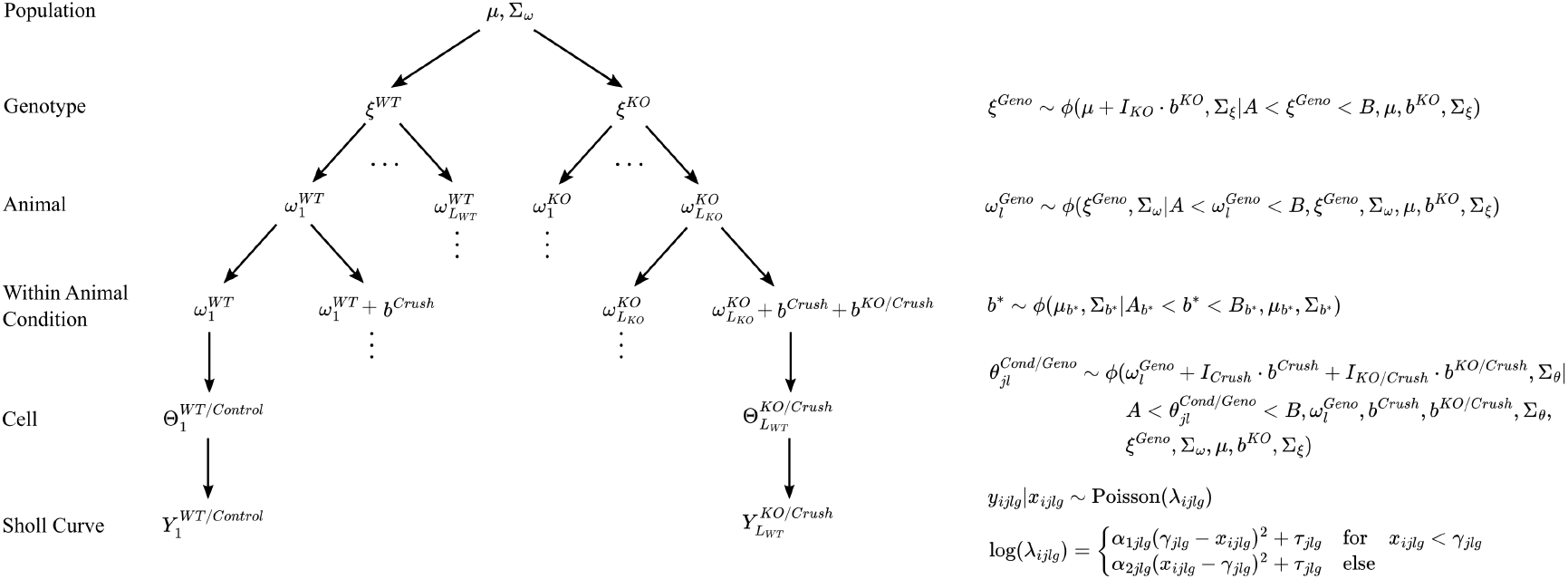
Hierarchical structure for model 3. As before, all notation is shared with models displayed in Figures 2.3 and 2.4, except *ξ* denotes genotype-level parameters. Additionally, *b^KO^* denotes the genotype level effect, *b^Crush^* denotes the condition level effect, and *b^KO/Crush^* denotes the interaction effect between condition and genotype. *I_*_* is an indicator variable equal to 1 for observations in group *, and 0 else.

The effect is given by

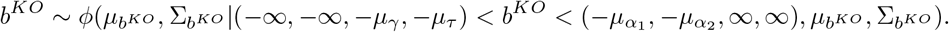

In our real data example, cell-level curves correspond to either a control eye, or an eye subject to optical nerve crush injury. Each animal has a crush and control eye, so we model the effect of condition, and the interaction of condition and genotype at the cell level via

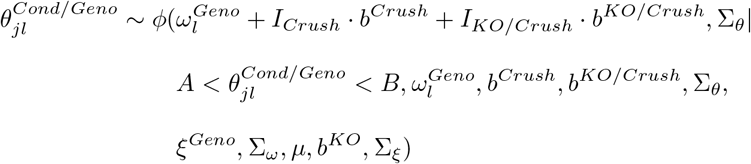

where

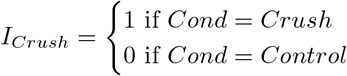

and

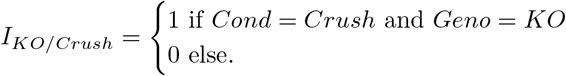

The effect of condition is given by

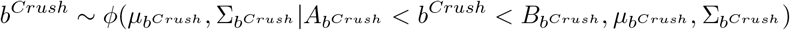

where

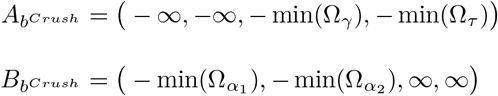

and 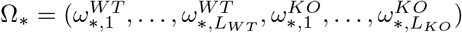.

The interaction effect is

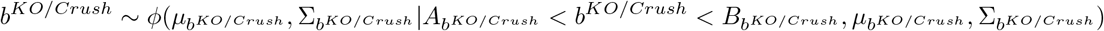

where

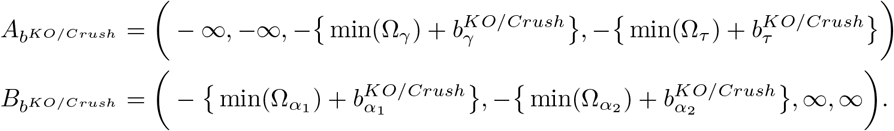

As before, we assume half-t priors on all standard deviation parameters.

All models were fit using MCMC via rjags (version 4-13).

### 2.6 Simulation Study

We limit simulation to model 2 (Section 2.4), except we incorporate cell-level data in the model hierarchy. We simulate data under six scenarios, primarily considering changes for effects on *τ* at the group level. Denoting the two grouping variables as either condition or side, the assumed effects in each scenario are shown in Table 1.

**Table 1.** Effects for each simulation scenario. We only consider effects on *τ* as it’s the most relevant parameter for the purposes of this article.

| Scenario | Parameter | Effect |  |  |
| --- | --- | --- | --- | --- |
|  |  | Condition | Side | Interaction |
| 1 | $\alpha_1$ | 0.00 | 0.00 | 0.00 |
| | $\alpha_2$ | 0.00 | 0.00 | 0.00 |
| | $\gamma$ | 0.00 | 0.00 | 0.00 |
| | $\tau$ | 0.00 | 0.00 | 0.00 |
| 2 | $\alpha_1$ | 0.00 | 0.00 | 0.00 |
| | $\alpha_2$ | 0.00 | 0.00 | 0.00 |
| | $\gamma$ | 0.00 | 0.00 | 0.00 |
| | $\tau$ | 0.50 | 0.00 | 0.00 |
| 3 | $\alpha_1$ | 0.00 | 0.00 | 0.00 |
| | $\alpha_2$ | 0.00 | 0.00 | 0.00 |
| | $\gamma$ | 0.00 | 0.00 | 0.00 |
| | $\tau$ | 0.00 | -0.25 | 0.00 |
| 4 | $\alpha_1$ | 0.00 | 0.00 | 0.00 |
| | $\alpha_2$ | 0.00 | 0.00 | 0.00 |
| | $\gamma$ | 0.00 | 0.00 | 0.00 |
| | $\tau$ | 0.50 | -0.25 | 0.50 |
| 5 | $\alpha_1$ | 0.00 | 0.00 | 0.00 |
| | $\alpha_2$ | 0.00 | 0.00 | 0.00 |
| | $\gamma$ | 0.00 | 0.00 | 0.00 |
| | $\tau$ | 0.50 | -0.25 | 0.50 |
| 6 | $\alpha_1$ | 0.00 | 0.00 | 0.00 |
| | $\alpha_2$ | 0.00 | 0.00 | 0.00 |
| | $\gamma$ | 0.00 | 0.00 | 0.00 |
| | $\tau$ | 0.50 | -0.25 | 0.50 |

Unless otherwise indicated, all simulation parameters are identical to the baseline scenario. At baseline, we simulate data using 5 animals per group and 10 cells per animal. For scenario 5, we double the cells per animal assumed at baseline. The baseline group-level variance parameters are set as 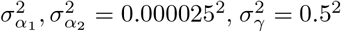, and 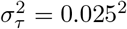, while baseline variance parameters for all other levels are set as 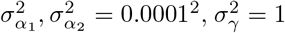, and 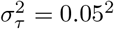. For scenario 6, we set 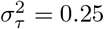 at the animal-level. Population level parameters are initialized as *μ* = (−0.002, −0.002, 30, 2) for each scenario.

To benchmark the proposed method, we first define the posterior probability that some effect *b* is less than 0 for simulation run *i* as *P_i_*(*b* < 0). Now define

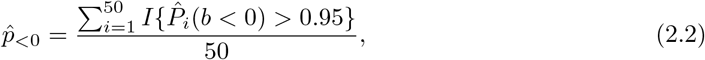

which is the proportion of simulation runs where the estimated posterior probability of some effect *b* having a negative sign is greater than 0.95. We estimate 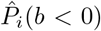 as the proportion of MCMC samples for parameter *b* that fall below 0. We can similarly define *P_i_*(*b* > 0) and

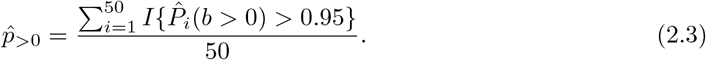

Across simulation runs, we expect 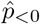 (or 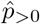) to approach 1 as the relative strength of a negative (or positive) effect increases. Similarly, if there is no true effect, we expect both 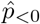 and 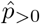 to be approximately 0. Similar to the estimated power and FPR in frequentist simulations, 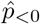 and 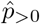 can be interpreted as the proportion of simulation runs where our decision criteria is met. Thus, we can add 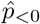 and 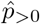 to measure the probability of a false discovery when there is no true effect.

## 3. Results/Application

### 3.1 Ungrouped Mouse Dataset

Investigators were interested in being able to assay changes in microglial morphology. To investigate this, sections of mouse cortical tissue were generated and underwent histology for a microglia-specific marker. Images of microglia in the primary visual cortex were collected, and Sholl analysis was performed to assay the number of microglial processes at regular distance intervals from the cell soma. These data were plotted as Sholl curves to represent the overall morphological profile of individual microglia. Original analysis detected a range of Sholl curve profiles at the level of individual cells which were used to generate animal level aggregate Sholl curves.

Figure 5 shows the fitted curves at each level of the model hierarchy. In this example, we are primarily interested in the model’s ability to capture the possible range of Sholl curves at any level of the hierarchy. This desired flexibility is particularly apparent at the cell level, where Sholl curves can vary greatly within an animal. For example, we see the model has no issues capturing the curve with abnormally large branch maximum associated with Gazer. There is not much variation between images within an animal, which isn’t surprising because images are taken of adjacent areas in the same brain region. The model is able to capture an overall animal level curve quite well, while also allowing for natural variations between animals.

**Fig. 5.**
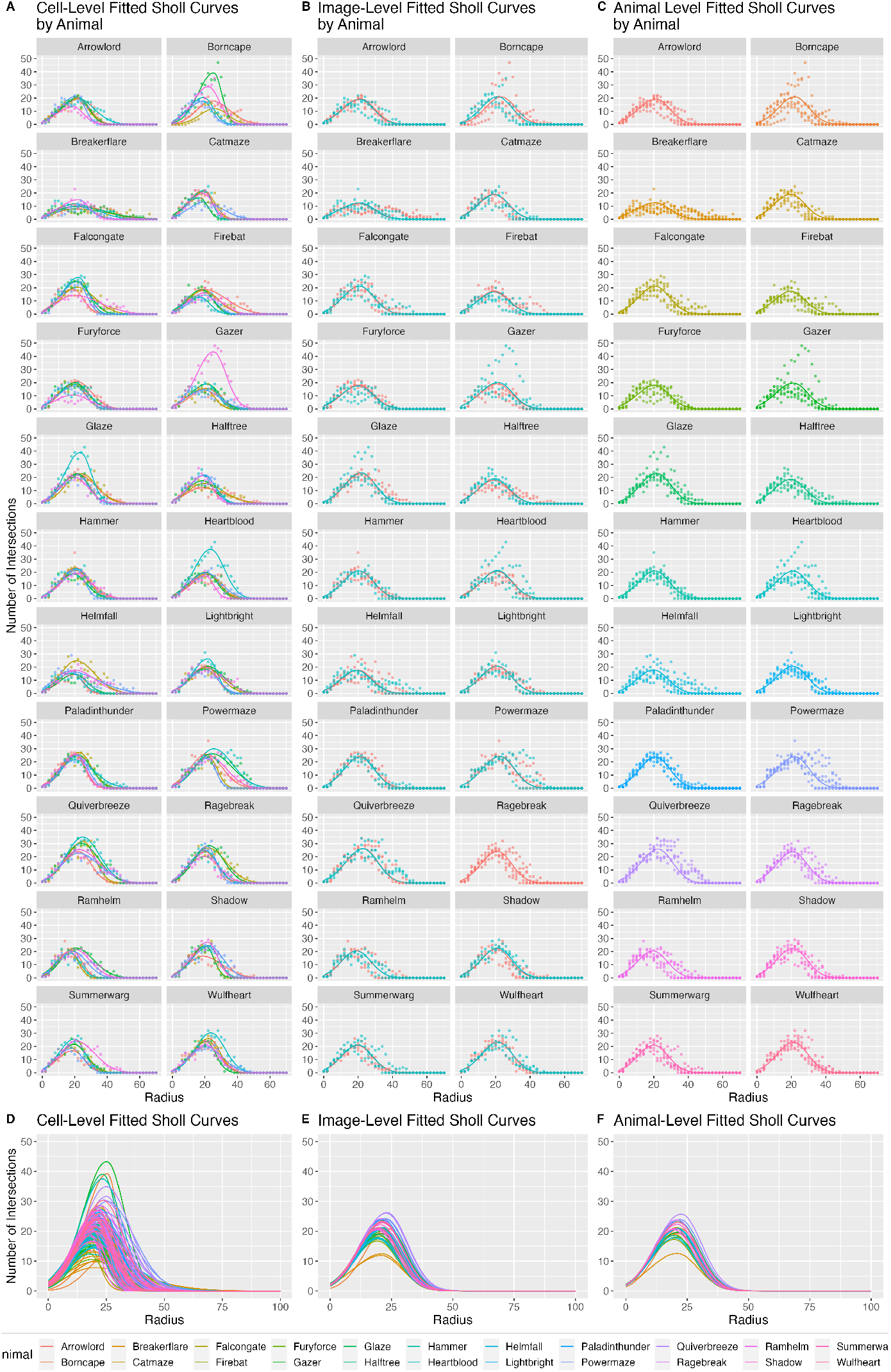
Fitted curves at each level of the model hierarchy, obtained with model 1 via MCMC. **A**: Cell-level fitted curves for each animal, where color indicates the cell. **B**: Image-level fitted curves for each animal, where color indicated image. **C**: Animal-level fitted curves. **D**: All cell-level fitted curves displayed in panel A, superimposed to show the cell-level variation. **E**: All image-level fitted curves displayed in panel B, superimposed to show the image-level variation. **F**: All animal-level fitted curves displayed in panel C, superimposed to show the animal-level variation.

### 3.2 MD/ND Dataset

The MD/ND dataset is one part of the data supporting findings from a study interested in the role of microglia in experience-dependent synaptic plasticity (Sipe *and others*, 2016). These data were used to demonstrate that ocular dominance plasticity induces hyper-ramification of microglia and that this effect is limited to the cortical area undergoing plasticity, which is the contralateral binocular visual cortex.

To examine whether microglia play a role in the process of experience-dependent synaptic plasticity, investigators assessed microglial response by assaying changes in microglial morphology after inducing ocular dominance plasticity. Tissue sections were generated from wildtype mice that had been monocularly deprived via eyelid suturing for 12 hours. Sections underwent histology for a microglia-specific marker and images of the binocular primary visual cortex were generated in both brain hemispheres to include visual areas both contralateral and ipsilateral to the deprived eye. Sholl analysis was performed on individual microglia. Analysis of these data were performed by constructing animal level aggregate Sholl curves, and fitting an ANOVA at each radius of the aggregate curves, which were used to test differences in process crossings between experimental conditions in both cortical hemispheres.

Though these data originally did contain multiple images and cells per animal, we re-analyze the truncated data using the proposed method. This allows both an example of how our method can be used when only animal level data is available, along with easier comparison with the original analysis. These data contain two grouping variables: condition and side. Condition is either monocular deprivation (MD) or no deprivation (ND), and side is either ipsilateral (I) or contralateral (C).

The fitted curves are displayed in Figure 6, which, as seen in panel A, do well at capturing the Sholl curves. In panel B we see the curve for group MD/C is quite large relative to other groups, indicating potential hyper-ramification of cells in this group. We show 95% credible intervals for group and interaction effects in Figure 7. These are superimposed over the approximate posterior density obtained via MCMC. Most parameters have posterior mass roughly centered at 0, with the exception of interaction effects on τ and *γ*. Clearly most of the posterior mass for both these effects falls above 0, which indicates a positive interaction effect associated with these parameters. To perform formal inference, we define a cutoff of 0.95, and check if an effect has at least 95% of its posterior mass either above or below 0. Table 2 shows the estimated posterior probability each effect is less than 0. We say an effect exists in the positive direction for a parameter if this estimate is less than 0.05. Conversely, an effect exists in the negative direction if this estimate is more than 0.95. With respect to this cutoff, we can say a positive interaction effect exists for both τ and γ, while no other effects succeed in meeting this criteria. Recall the branch maximum and critical value are given by *e^τ^* and γ in our model, respectively. This suggests microglia hyperramification is indeed limited to the contraleteral binocular visual cortex, which agrees with the original analysis of these data.

**Table 2.** Estimated posterior probability of a negative effect for each parameter in the MD/ND model. Quantities are estimated as the proportion of MCMC samples that fall below 0.

| Parameter | $\hat{P}(\text{Effect} < 0)$ | | |
| --- | --- | --- | --- |
|  | Condition | Side | Interaction |
| $\alpha_1$ | 0.41 | 0.39 | 0.33 |
| $\alpha_2$ | 0.49 | 0.46 | 0.32 |
| $\gamma$ | 0.60 | 0.85 | 0.02 |
| $\tau$ | 0.68 | 0.87 | 0.00 |

**Fig. 6.**
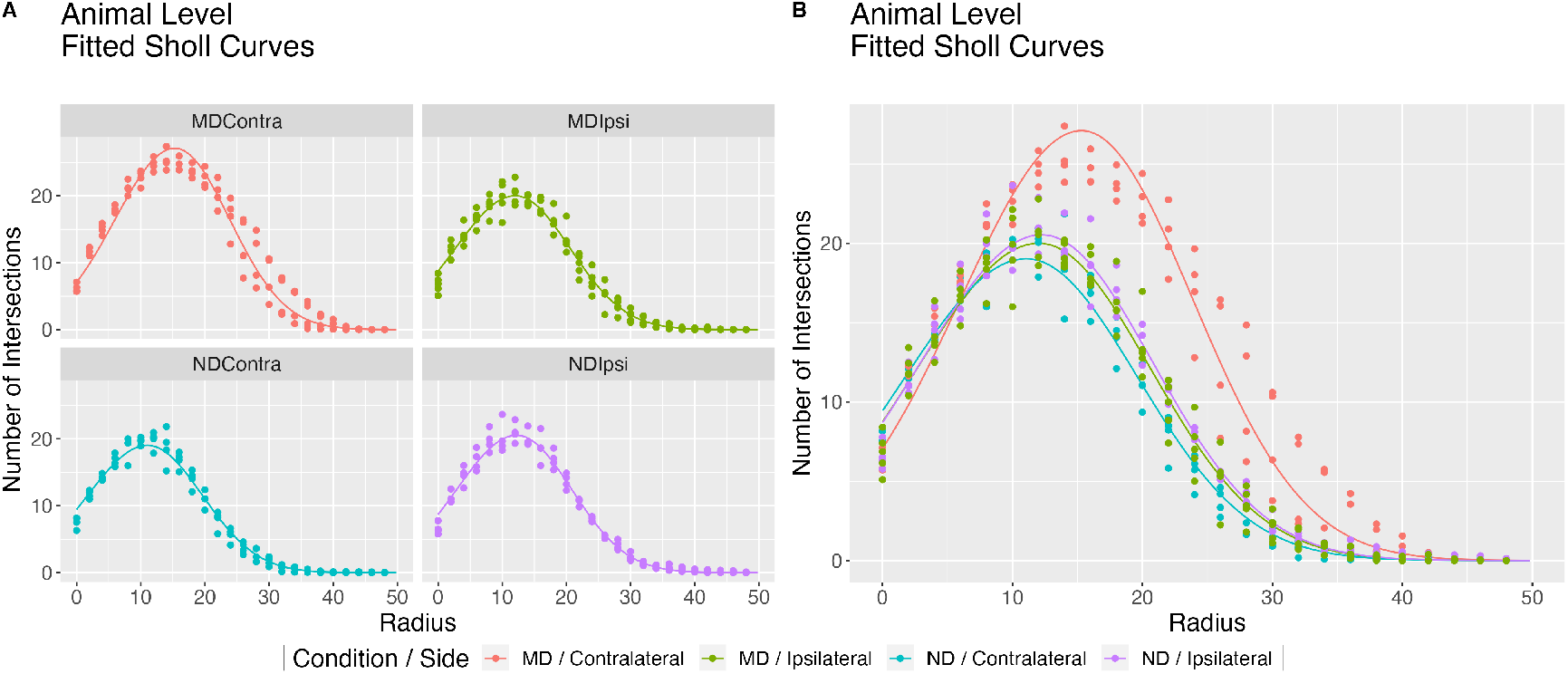
Group-level fitted curves obtained by fitting model 2 to the MD/ND dataset. **A**: Fitted curves faceted by group, superimposed over animal level Sholl curves. **B**: All four facets from panel A, super-imposed to better show hyper-ramification of the MD/Contra group.

**Fig. 7.**
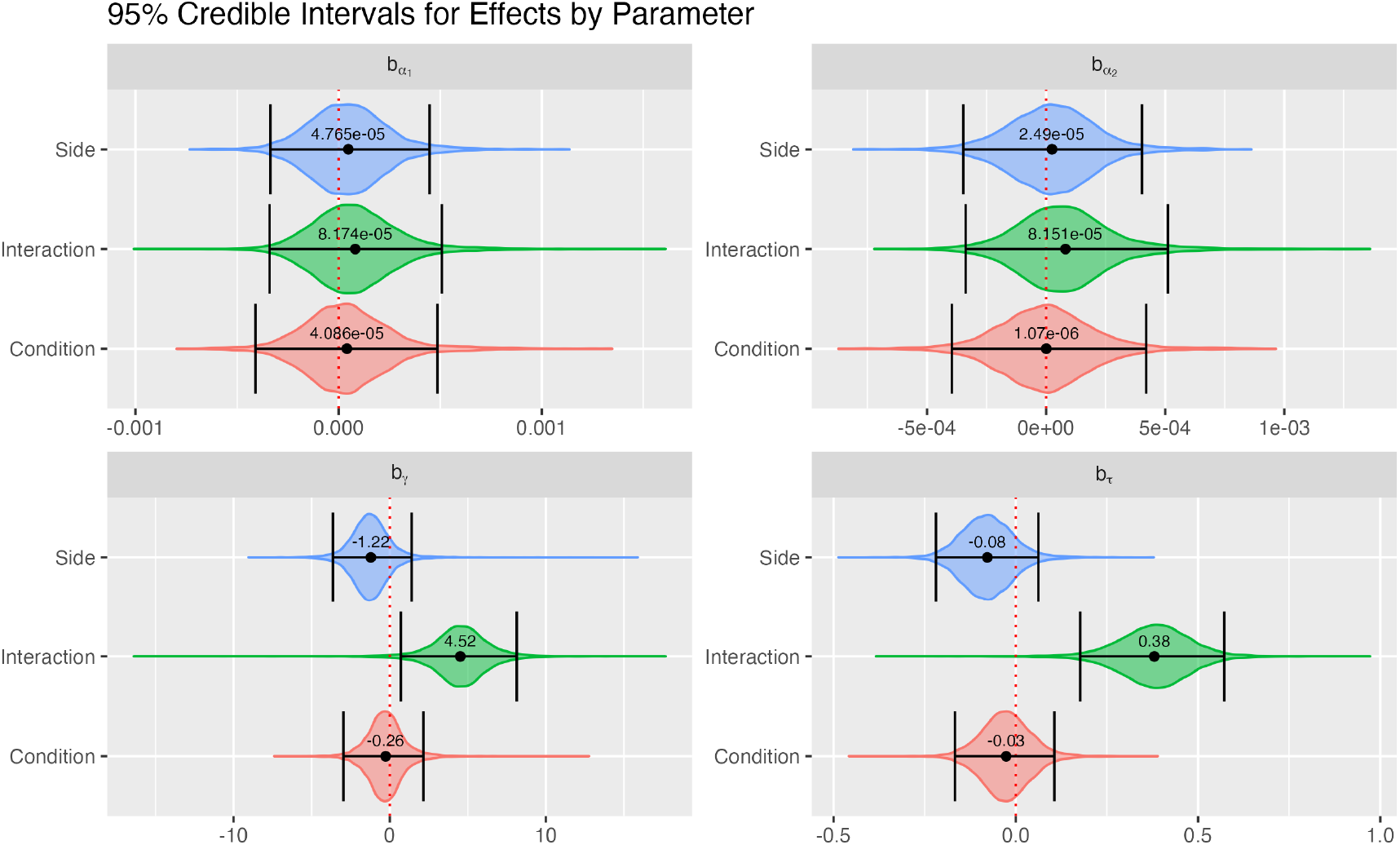
95% credible intervals for each effect in model 2, fitted to the MD/ND dataset. Credible intervals are computed as the highest density posterior interval. Credible intervals are superimposed over the approximate posterior distributions obtained via MCMC. Estimated posterior means are represented by black dots with point estimates displayed above. The dotted red line is fixed at 0.

### 3.3 GPNMB Knockout Dataset

Investigators were interested in the effect of the loss of transmembrane glycoprotein NMB (GPNMB) on the microglial response to an optic nerve crush (ONC) injury. GPNMB can work to reduce inflammation and is highly expressed in microglia, so the presence or absence of GPNMB may influence the role of microglia in the retina following ONC injury. An ONC injury was performed on a pilot cohort of 9 mice. Mice had either wildtype expression of GPNMB or a genetic knockout. For each animal, crush was performed on one eye and the contralateral eye underwent a sham injury which served as an inter-animal control. Retinas were collected 7 days after injury. Retinas were stained for microglia-specific markers and the ganglion cell layer/inner plexiform layer was imaged using confocal microscopy. Sholl analysis was performed on individual microglia and, similar to above, analysis was performed by constructing animal level aggregate Sholl curves. An ANOVA with repeated measures was conducted on both the branch maximum and critical value to test the differences between genotype, condition, and interaction. Results of this analysis are shown in Table 3.

**Table 3.** Two-way ANOVA with repeated measures fit to the branch maximum and critical value of the truncated GPNMB knockout data

| Response | Effect | nDF | dDF | F | p-value |
| --- | --- | --- | --- | --- | --- |
| Branch Maximum | Genotype | 1 | 5 | 14.05 | 0.01 |
|  | Condition | 1 | 5 | 34.62 | 0.00 |
|  | Interaction | 1 | 5 | 10.89 | 0.02 |
| Critical Value | Genotype | 1 | 5 | 2.66 | 0.16 |
|  | Condition | 1 | 5 | 14.12 | 0.01 |
|  | Interaction | 1 | 5 | 0.23 | 0.65 |

Cell-level fitted curves, separated by animal are displayed in Figure 8. Figure 9 shows 95% credible intervals for effects on genotype, condition, and interaction, superimposed over approximate posterior distributions. As with the MD/ND example, we report the estimated posterior probability each effect is less than 0 in Table 4. Using an 0.95 cutoff as before, we see the proposed method and two-way ANOVA detected similar effects on the branch maximum, while the proposed method also detects genotype and interaction effects on the critical value. Additionally, our method offers increased granularity when differences between curves are not obvious. Unlike the MD/ND example, visual differences between fitted curves in Figure 8 are not limited to these two summaries. Using our method, we are able to quantify these differences by leveraging *α*_1_ and *α*_2_, rather than only relying on the curve maximum. Specifically, we detect a negative condition effect on *α*_1_, meaning crush curves have steeper growth states than control curves. Though we only report effects here, we also have the option to investigate parameters, associated variance terms, and combinations of parameters (such as the y-intercept) at each level of the model hierarchy, providing a rich toolbox for investigating subtle curve differences.

**Table 4.** Estimated posterior probability of a negative effect for each parameter in the GPNMB knockout model. Quantities are estimated as the proportion of MCMC samples that fall below 0.

| Parameter | $\hat{P}(\text{Effect} < 0)$ | | |
| --- | --- | --- | --- |
|  | Condition | Genotype | Interaction |
| $\alpha_1$ | 1.00 | 0.02 | 0.44 |
| $\alpha_2$ | 0.89 | 0.32 | 0.40 |
| $\gamma$ | 1.00 | 0.00 | 1.00 |
| $\tau$ | 0.93 | 0.01 | 1.00 |

**Fig. 8.**
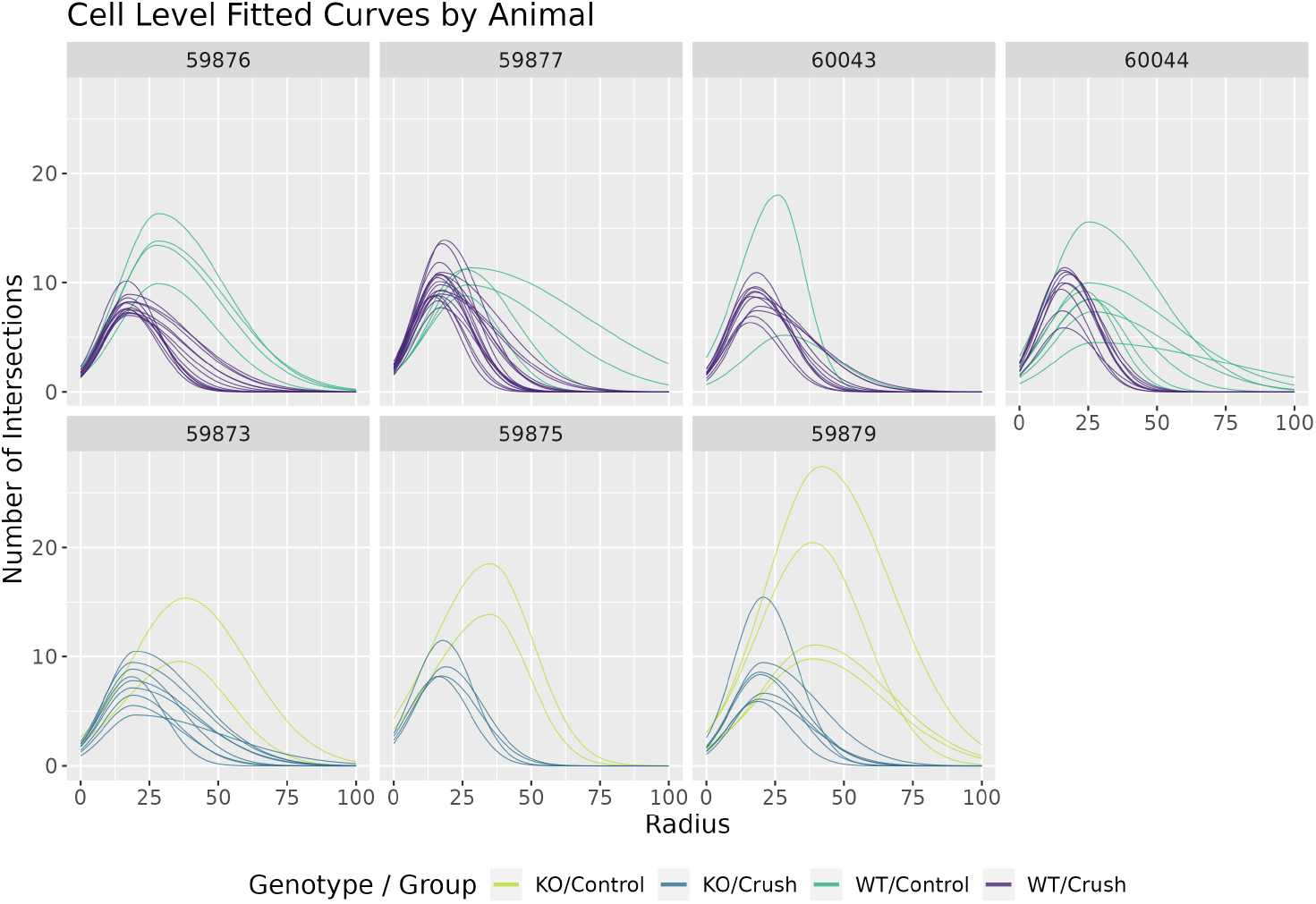
Cell-level fitted curves faceted by animal obtained by fitting model 3 to the GPNMB knockout dataset. An animal is either wild-type (WT), or has gene GPNMB knocked out (KO). Cells are associated with either an eye subject to optical nerve crush injury, or control. Each animal has both a crush eye and a control eye.

**Fig. 9.**
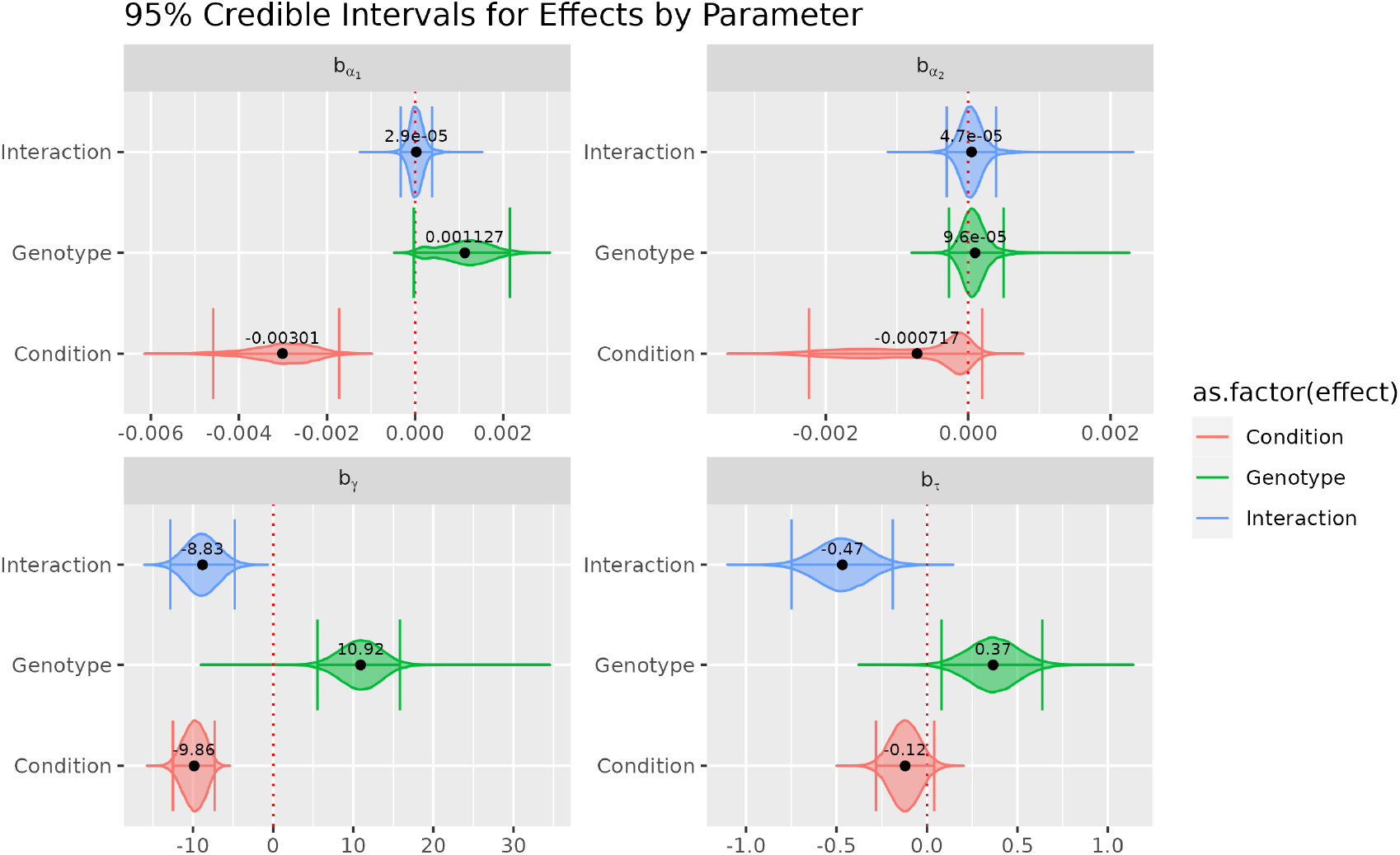
95% credible intervals for each effect in model 3, fitted to the GPNMB knockout dataset. Credible intervals are computed as the highest density posterior interval. Credible intervals are superimposed over the approximate posterior distributions obtained via MCMC. Estimated posterior means are represented by black dots with point estimates displayed above. The dotted red line is fixed at 0.

## 4. Simulation Study

We compare the proposed method to the existing ANOVA based method discussed in Section 2.1 via simulation study. We narrow the scope of simulation to model 2 (Section 2.4), i.e. the simplest model with effects, but we include an additional level of hierarchy for cell-level data. The simulated data is truncated at the animal level in order to apply ANOVA, while the full data is used in the proposed model.

We simulate data under the assumed hierarchical structure of the proposed model. We only consider effects on *τ* since the critical value is both a popular summary metric and the most relevant parameter to the examples we discussed in Section 3. Data are simulated under six scenarios: 1) baseline scenario with no effects, 2) moderate positive marginal effect, 3) small negative marginal effect, 4) both effects in the previous two scenarios with moderate positive interaction effect, 5) identical to scenario 4 except twice as many cells per animal, 6) identical to scenario 4 except with large animal-level variance.

50 datasets are simulated for each scenario, and both models are fit to each dataset. Details regarding the MCMC sampling procedure and diagnostics are included in the Supplementary Materials. In Table 5, we report the frequentist power and FPR estimates for the ANOVA at the 0.05 level. Also reported in Table 5 are 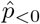 and 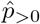 for the proposed method.

**Table 5.** Estimated type-I error rate (FPR) and power for ANOVA, compared with 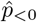 and 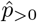 for the proposed model under each simulation scenario. Power and FPR for the ANOVA model are estimated at the 0.05 level of significance. 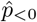 and 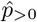 for the proposed model are computed via Equations 2.2 and 2.3. Dashes fill cells where there is no relevant quantity.

| Scenario | Parameter | Method |  |  |  |  |  |  |  |  |  |  |  |
| --- | --- | --- | --- | --- | --- | --- | --- | --- | --- | --- | --- | --- | --- |
|  |  | Proposed |  |  |  |  |  | ANOVA |  |  |  |  |  |
|  |  | Condition |  | Side |  | Interaction |  | Condition |  | Side |  | Interaction |  |
| | | $\hat{p}_{<0}$ | $\hat{p}_{>0}$ | $\hat{p}_{<0}$ | $\hat{p}_{>0}$ | $\hat{p}_{<0}$ | $\hat{p}_{>0}$ | Power | FPR | Power | FPR | Power | FPR |
| 1 | $\alpha_1$ | 0.04 | 0.00 | 0.00 | 0.02 | 0.00 | 0.00 | — | — | — | — | — | — |
| | $\alpha_2$ | 0.02 | 0.02 | 0.06 | 0.02 | 0.00 | 0.00 | — | — | — | — | — | — |
| | $\gamma$ | 0.04 | 0.08 | 0.02 | 0.02 | 0.04 | 0.00 | — | 0.06 | — | 0.06 | — | 0.06 |
| | $\tau$ | 0.10 | 0.02 | 0.08 | 0.06 | 0.04 | 0.14 | — | 0.12 | — | 0.12 | — | 0.16 |
| 2 | $\alpha_1$ | 0.00 | 0.02 | 0.00 | 0.00 | 0.00 | 0.00 | — | — | — | — | — | — |
| | $\alpha_2$ | 0.02 | 0.02 | 0.02 | 0.00 | 0.00 | 0.00 | — | — | — | — | — | — |
| | $\gamma$ | 0.04 | 0.02 | 0.02 | 0.00 | 0.00 | 0.00 | — | 0.06 | — | 0.08 | — | 0.06 |
| | $\tau$ | 0.00 | 1.00 | 0.04 | 0.08 | 0.08 | 0.06 | 1.00 | — | — | 0.18 | — | 0.14 |
| 3 | $\alpha_1$ | 0.02 | 0.00 | 0.00 | 0.02 | 0.00 | 0.00 | — | — | — | — | — | — |
| | $\alpha_2$ | 0.02 | 0.02 | 0.00 | 0.00 | 0.00 | 0.00 | — | — | — | — | — | — |
| | $\gamma$ | 0.02 | 0.08 | 0.00 | 0.04 | 0.00 | 0.02 | — | 0.04 | — | 0.10 | — | 0.04 |
| | $\tau$ | 0.04 | 0.18 | 0.92 | 0.00 | 0.10 | 0.06 | — | 0.22 | 0.94 | — | — | 0.18 |
| 4 | $\alpha_1$ | 0.00 | 0.02 | 0.00 | 0.00 | 0.00 | 0.02 | — | — | — | — | — | — |
| | $\alpha_2$ | 0.02 | 0.00 | 0.00 | 0.00 | 0.00 | 0.00 | — | — | — | — | — | — |
| | $\gamma$ | 0.00 | 0.02 | 0.00 | 0.02 | 0.02 | 0.02 | — | 0.02 | — | 0.08 | — | 0.00 |
| | $\tau$ | 0.00 | 1.00 | 0.80 | 0.00 | 0.00 | 1.00 | 1.00 | — | 0.56 | — | 0.98 | — |
| 5 | $\alpha_1$ | 0.02 | 0.00 | 0.00 | 0.00 | 0.00 | 0.00 | — | — | — | — | — | — |
| | $\alpha_2$ | 0.02 | 0.00 | 0.00 | 0.02 | 0.00 | 0.00 | — | — | — | — | — | — |
| | $\gamma$ | 0.02 | 0.02 | 0.00 | 0.08 | 0.00 | 0.02 | — | 0.06 | — | 0.02 | — | 0.12 |
| | $\tau$ | 0.00 | 1.00 | 0.94 | 0.00 | 0.00 | 1.00 | 1.00 | — | 0.34 | — | 0.94 | — |
| 6 | $\alpha_1$ | 0.00 | 0.00 | 0.00 | 0.00 | 0.00 | 0.00 | — | — | — | — | — | — |
| | $\alpha_2$ | 0.00 | 0.04 | 0.02 | 0.02 | 0.00 | 0.00 | — | — | — | — | — | — |
| | $\gamma$ | 0.02 | 0.08 | 0.00 | 0.08 | 0.04 | 0.02 | — | 0.06 | — | 0.02 | — | 0.12 |
| | $\tau$ | 0.00 | 0.96 | 0.38 | 0.00 | 0.00 | 0.72 | 1.00 | — | 0.14 | — | 0.44 | — |

In scenario 1, when there is no true effect, both methods perform comparably. For the proposed method, when the larger true effect exists for condition and/or interaction, 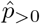 is almost always 1. For the smaller effect on side, we see slightly smaller values for 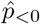, which are approximately 0.92 and 0.80 in scenarios 3 and 4, respectively. In scenario 5, when cells per animal is increased, we see 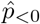 for side increase to 0.94. Our method seems to struggle when variance terms are large relative to the effect size. Specifically, in scenario 6, we see 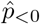 for side and 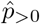 for interaction dip to 0.38 and 0.72, respectively. In contrast, the ANOVA based method, while fully powered in some scenarios, struggles when more effects and data are introduced. This is apparent in scenario 4, where ANOVA struggles to detect the side effect in the presence of both condition and interaction effects. Additionally, when the cells per animal is increased in scenario 5, we actually see a decrease in power to detect the side effect relative to scenario 4. In scenario 6, ANOVA reports a power of 0.14 for detecting the side effect, which is not much larger than the FPR of 0.12 when no effect is present. Overall, the proposed method performs favorably across scenarios when true effects are present, while, at worse, performing comparably to ANOVA at controlling the false discovery rate.

## 5. Discussion

Sholl analysis is still the gold standard for morphological analysis in the microglia community. We propose a model for directly modeling Sholl curves, filling a long existing gap in the morphological inference pipeline. We generalize this model to a hierarchical Bayesian framework which naturally captures the nested structure of microglia imaging datasets. We apply our model in three real data examples and compare the proposed model to the analysis method previously applied to these data via simulation study.

Our applied examples showcase the flexibility of our method in capturing the myriad of shapes Sholl curves can take. We also demonstrate our model’s ability to capture relevant effects, potentially existing at multiple levels of the hierarchy. In our simulation study, we show the proposed method performs well when true effects are present, while being comparable to the competing method at controlling false discovery. In comparison, the ANOVA based method can be fully powered when a large enough effect exists but can become problematic as more data below the level of truncation is made available, in the presence of many true effects, or when the effect size is too small.

Our method can have some trouble sampling *α*_1_ and associate effects. Often we see low effective sample size relative to other parameters, along with difficulty getting chains to converge. That latter is reflected in both the Rhat and the approximate posterior for *α*_1_ effects in Figure 9. This could be alleviated by re-parameterizing the model. Instead of modeling the growth curve with *α*_1_, we can instead model the y-intercept directly. This may even be the preferred parameterization if estimating the number of processes originating from the soma is of interest. Our simulation study was also limited to effects on *τ*, leaving the door open for more rigorous study of other model parameters. Work can be done to relax assumptions on variance terms in the model, i.e. allowing for more than one shared variance parameter at each level of the hierarchy. Additionally, there are two primary ways this model can be further generalized: allowing for a negative binomial random component and including nonlinear parameters *κ*_1_ and *κ*_2_ in the mean model via

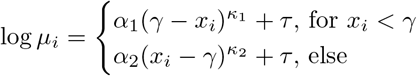

for *κ*_1_, *κ*_2_ > 1. The former relaxes the mean-variance relationship of the Poisson model, while the latter allows more flexible characterization of the growth and decay states.

In summary, we believe Sholl based morphological analyses can greatly benefit from modelbased methods which utilize all available data. Though the applied examples in this paper are limited to microglia, Sholl analysis is also a common method for quantifying the morphology of other cells, particularly neurons. We predict that our method is flexible enough to adequately capture the Sholl curve of other cell types, though modifications should be made to the specific model hierarchy to match the experimental design. We developed this method as a step toward more rigorous morphological analysis when Sholl analysis is the preferred method to quantify cell morphology. We anticipate that the proposed methodology will lead to improved analysis of microglial images by uncovering the changes in morphology that are most predictive of alterations in microglial function.

## 6. Supplementary Materials

The reader is referred to the online Supplementary Materials for technical appendices and annotated R code.

## Acknowledgments

We would like to thank Richard Libby along with the Libby lab for their data contributions.

## Funding

The project described in this publication was supported by the University of Rochester CTSA award number UL1TR002001 from the National Center for Advancing Translational Sciences of the National Institutes of Health. The content is solely the responsibility of the authors and does not necessarily represent the official views of the National Institutes of Health.

The data described in this publication was gathered by investigators supported by the National Institutes of Health (R01 NS114480, RO1 AA027111, T32 NS115705) and the SPIN foundation.

## Supplementary Material

**Fig. S1.**
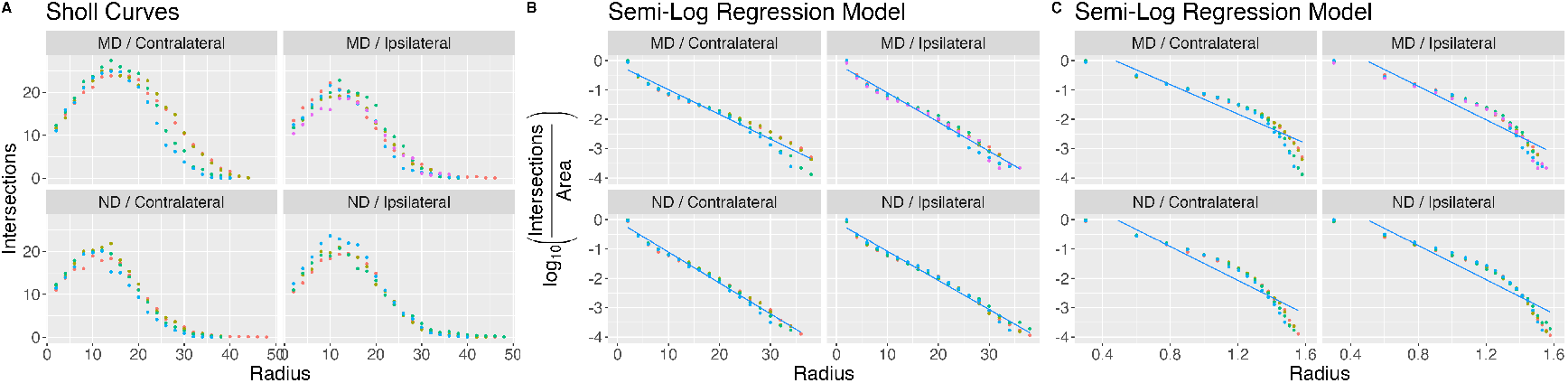
**A**: Aggregate Sholl curves from either the ipsilateral or contralateral side of mice subject to either monocular deprivation (MD) or control (ND). **B**: The semi-log model fit to the Sholl curves displayed in panel A. Notice how the transformed data oscillates about the fitted curve despite the linearization appearing adequate. Additionally, there appears to be mild heteroskedasticity near the tail of the transformed data. **C**: The log-log method fit to the Sholl curves displayed in panel A. The tail of the transformed curves drops quickly, suggesting this model may be more suited for Sholl curves with longer tails.

**Fig. S2.**
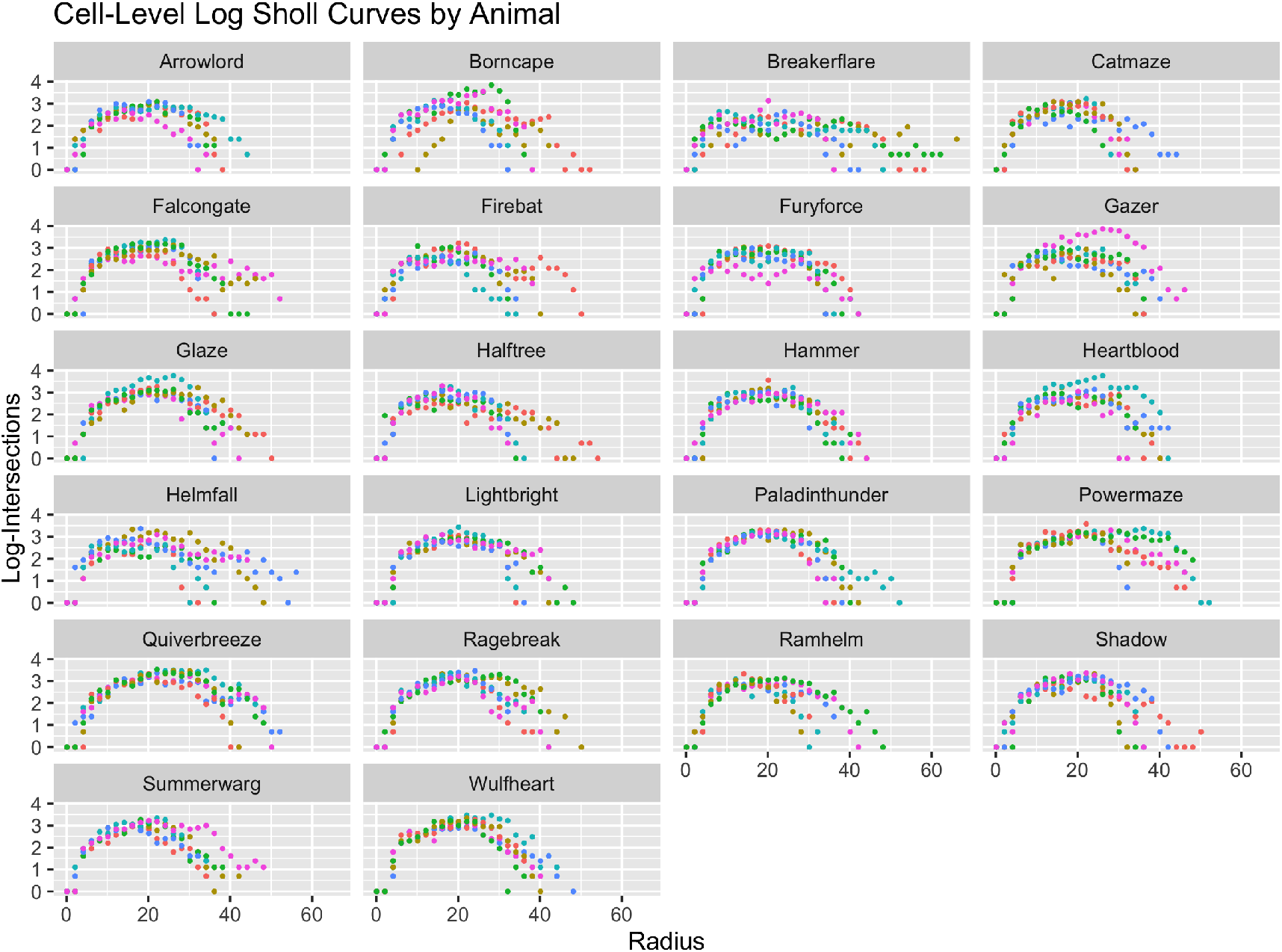
Cell-level log-transformed Sholl curves are displayed for the ungrouped animal dataset, where each facet corresponds to a difference animal. Our proposed model (Equation 2.1) fits a piece-wise parabola to the log-mean process crossings, which captures the structure of the data directly.

### S0.1 MCMC Sampling Procedures

#### S0.1.1 Ungrouped Mouse Dataset

We sample from the posterior using the JAGS implementation of MCMC, running 4 chains in parallel. For each chain, we we allow the sampler to adapt for 5000 iterations, followed by a 50000 iteration burn-in. The sampler is then run for 150000 iterations. Autocorrelation is alleviated by thinning, keeping a sample every 50 iterations.

#### S0.1.2 MD/ND Dataset

For each of 4 chains, we adapted a JAGS sampler for 5000 iterations, followed by a 50000 iteration burn in before running the sampler for 150000 iterations. Autocorrelation is alleviated by thinning, keeping a sample every 50 iterations.

#### S0.1.3 GPNMB Knockout Dataset

As before, the model is fit using a JAGS sampler and 4 chains, each of which are adapted with 10000 iterations. A burn-in of 250000 iterations was performed before obtaining 500000 samples. Auto-correlation is alleviated by thinning, keeping a sample every 50 iterations.

#### S0.1.4 Simulation Study

For each dataset, we simulate a seed for each of 4 chains which are all run in parallel. Then for each chain, a JAGS sampler was adapted for 5000 iterations, followed by a burn in of 15000 and 20000 iterations. Auto-correlation is alleviated by thinning, keeping a sample every 20 iterations.

### S0.2 Simulation Diagnostics

**Table S1.**
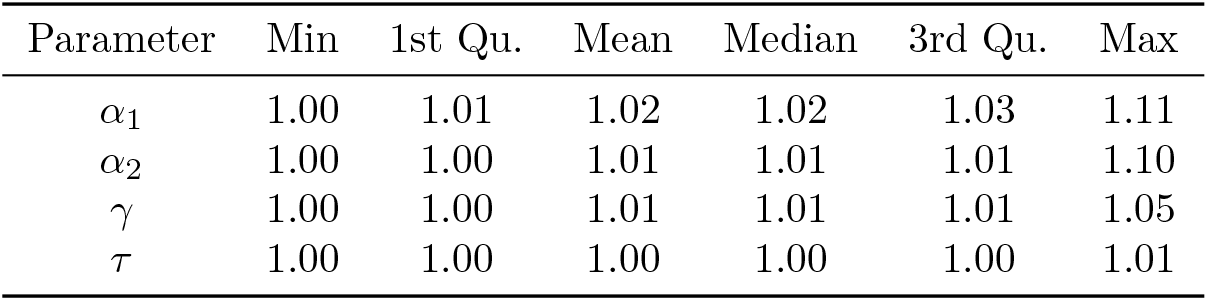
Summary of R-hat at the cell-level for simulation scenario 1. Due to the number of cell level parameters, the reported summaries are taken across the 50 simulation runs AND all cell-level parameters.

**Table S2.**
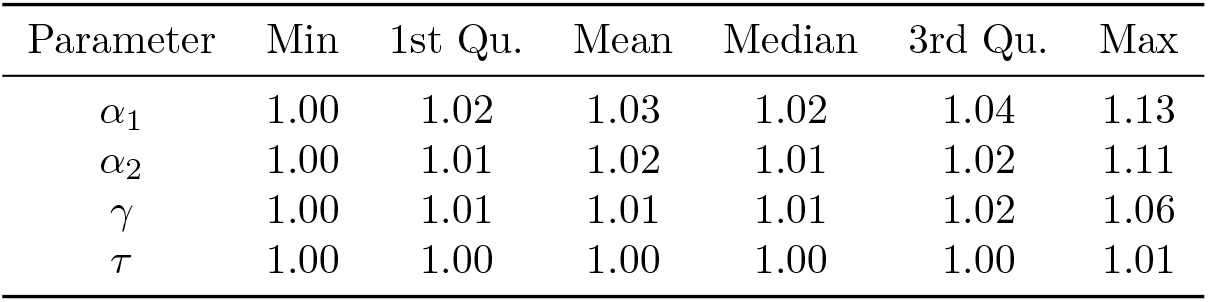
Summary of R-hat at the animal-level for simulation scenario 1. Due to the number of animal level parameters, the reported summaries are taken across the 50 simulation runs AND all animal-level parameters.

**Table S3.**
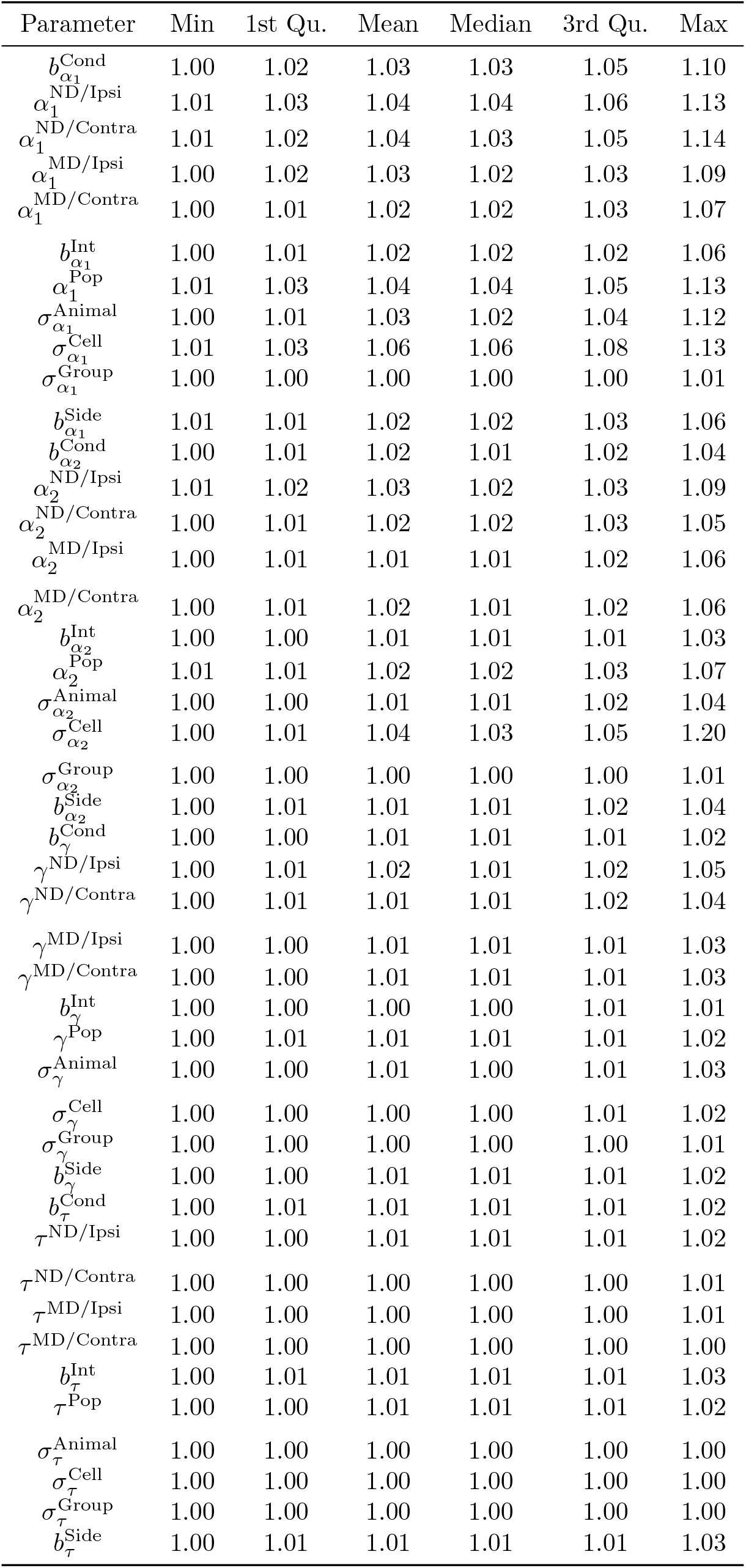
Summary of R-hat for other model parameters in simulation scenario 1. The reported summaries are taken across the 50 simulation runs.

**Table S4.**
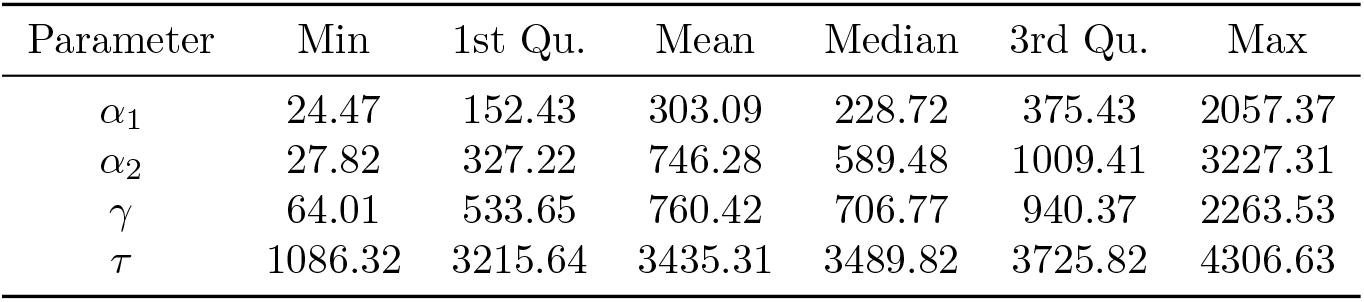
Summary of the effective sample size for parameters at the cell-level in simulation scenario 1. Due to the number of cell level parameters, the reported summaries are taken across the 50 simulation runs AND all cell-level parameters.

**Table S5.**
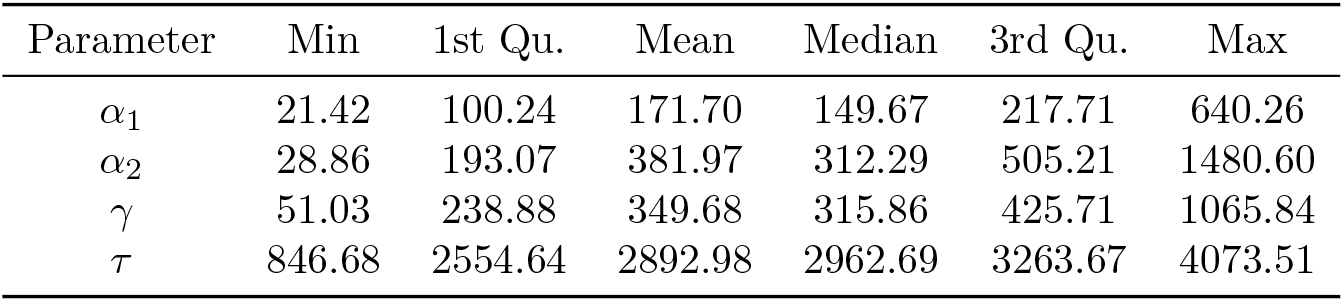
Summary of the effective sample size for parameters at the animal-level in simulation scenario 1. Due to the number of animal level parameters, the reported summaries are taken across the 50 simulation runs AND all animal-level parameters.

**Table S6.**
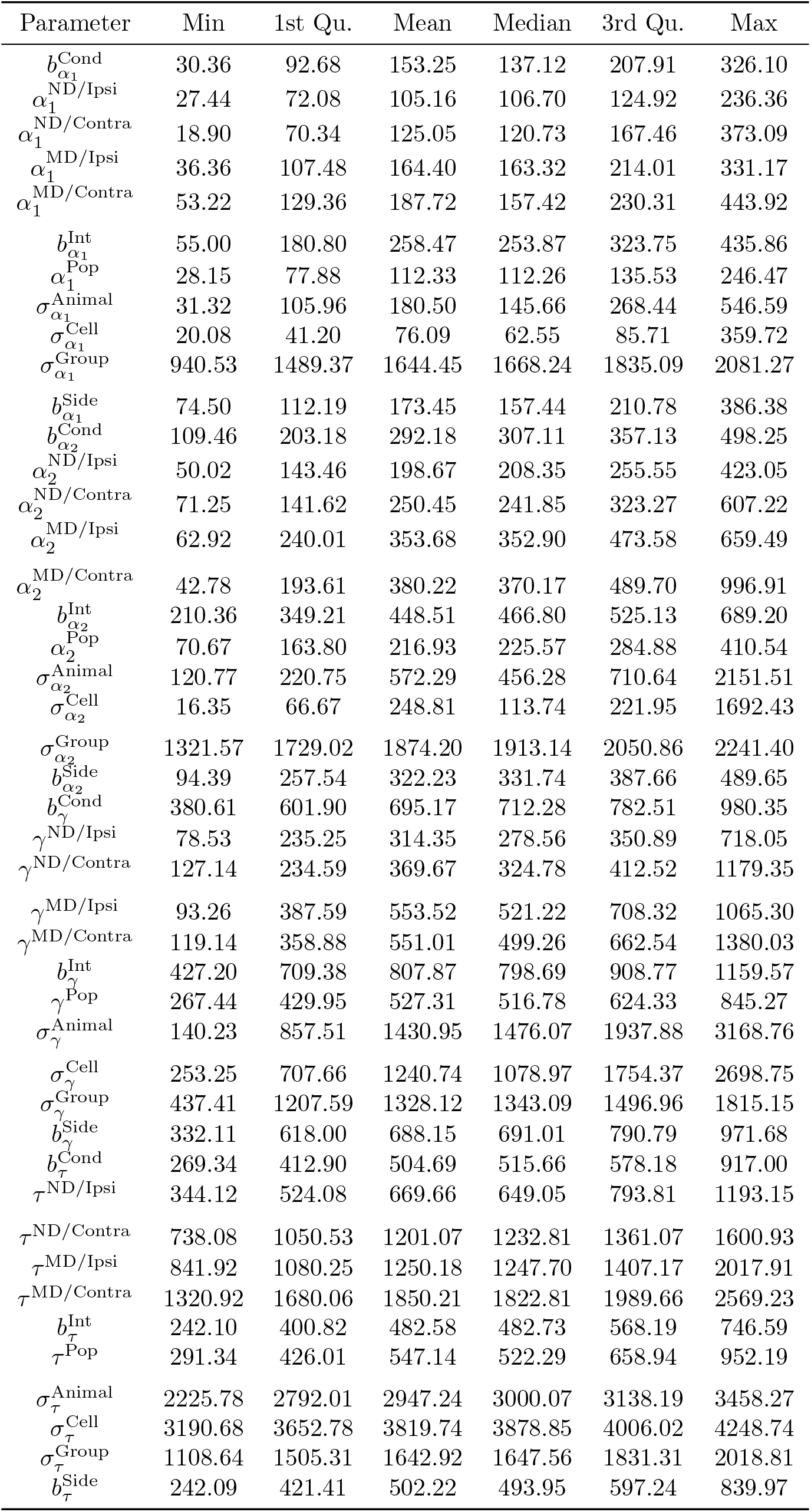
Summary of the effective sample size for parameters at the animal-level in simulation scenario 1. The reported summaries are taken across the 50 simulation runs.

**Table S7.**
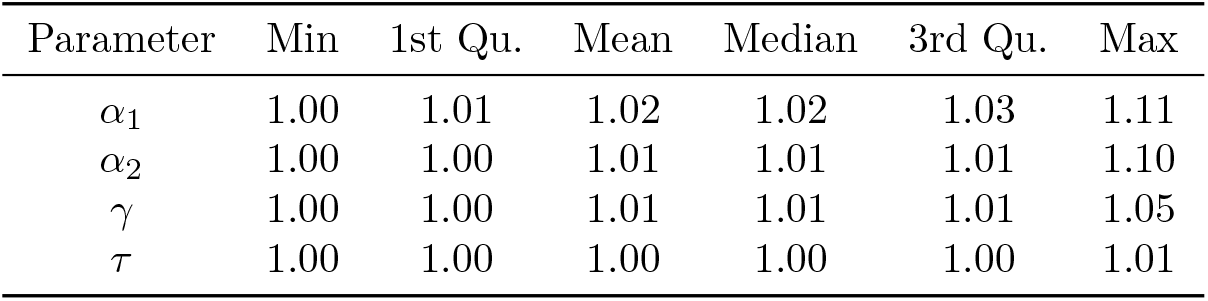
Summary of R-hat at the cell-level for simulation scenario 2. Due to the number of cell level parameters, the reported summaries are taken across the 50 simulation runs AND all cell-level parameters.

**Table S8.**
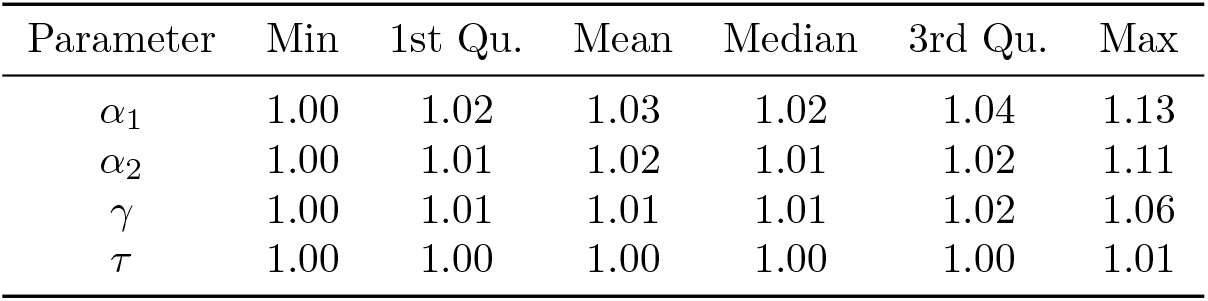
Summary of R-hat at the animal-level for simulation scenario 2. Due to the number of animal level parameters, the reported summaries are taken across the 50 simulation runs AND all animal-level parameters.

**Table S9.**
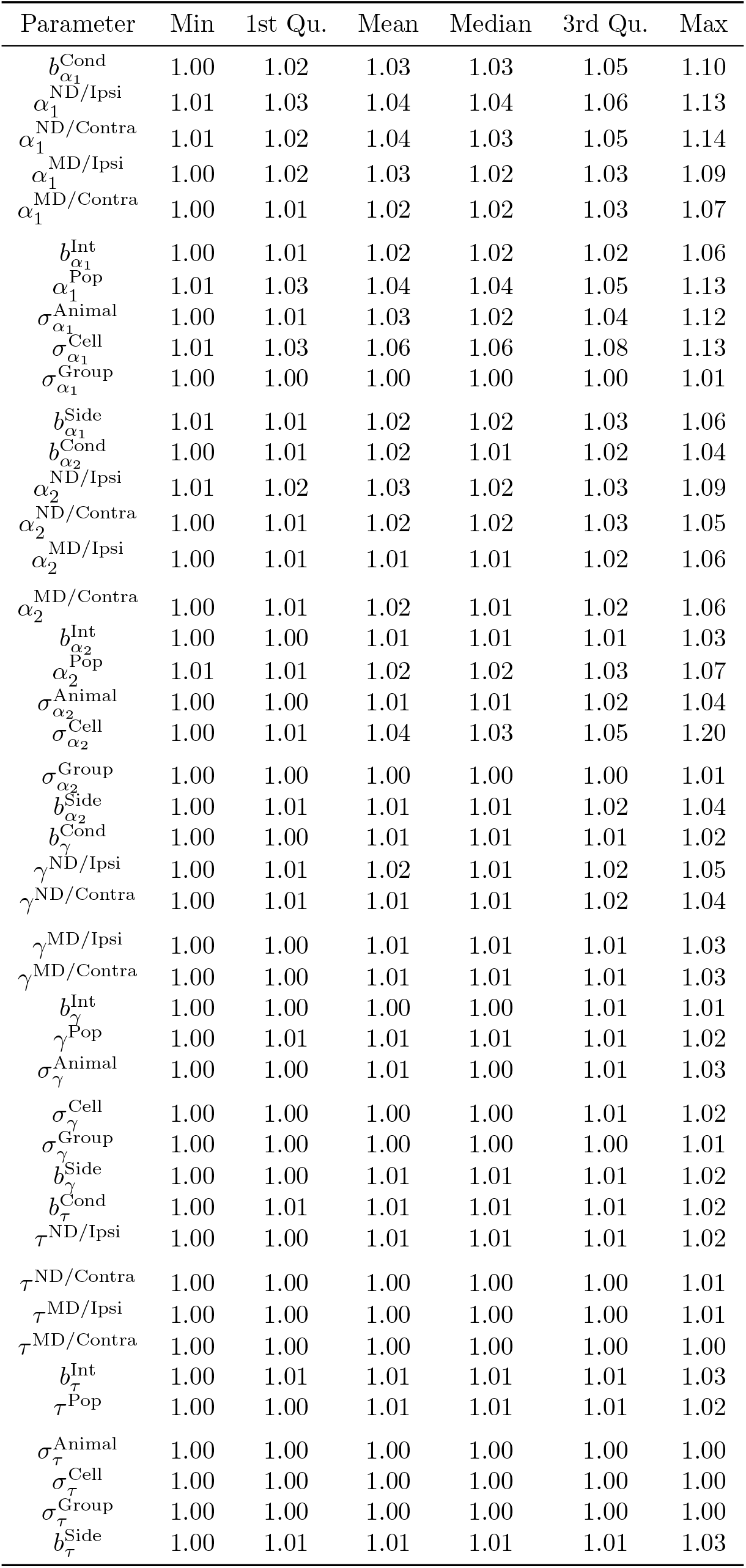
Summary of R-hat for other model parameters in simulation scenario 2. The reported summaries are taken across the 50 simulation runs.

**Table S10.**
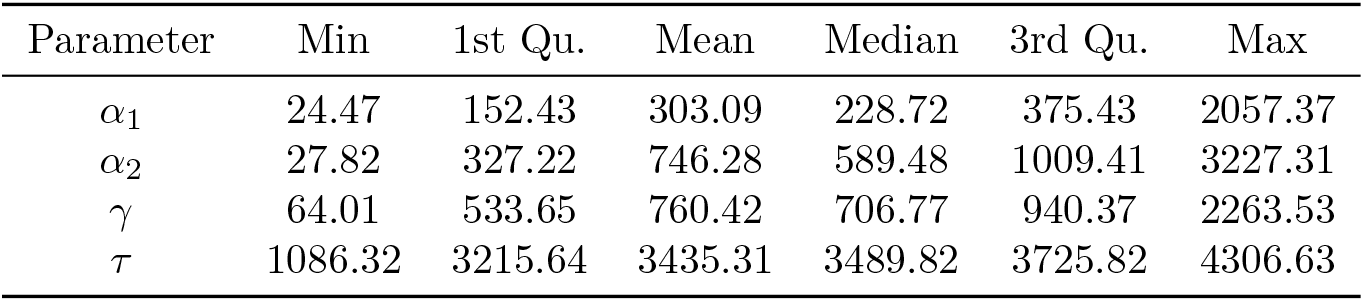
Summary of the effective sample size for parameters at the cell-level in simulation scenario 2. Due to the number of cell level parameters, the reported summaries are taken across the 50 simulation runs AND all cell-level parameters.

**Table S11.**
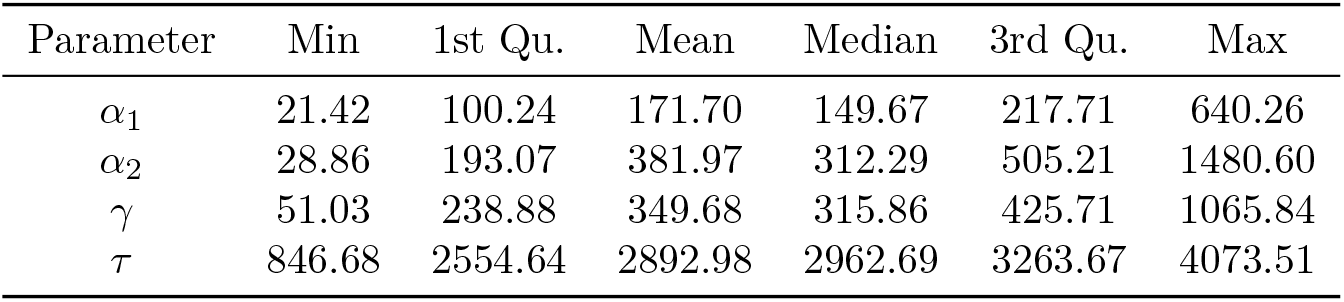
Summary of the effective sample size for parameters at the animal-level in simulation scenario 2. Due to the number of animal level parameters, the reported summaries are taken across the 50 simulation runs AND all animal-level parameters.

**Table S12.**
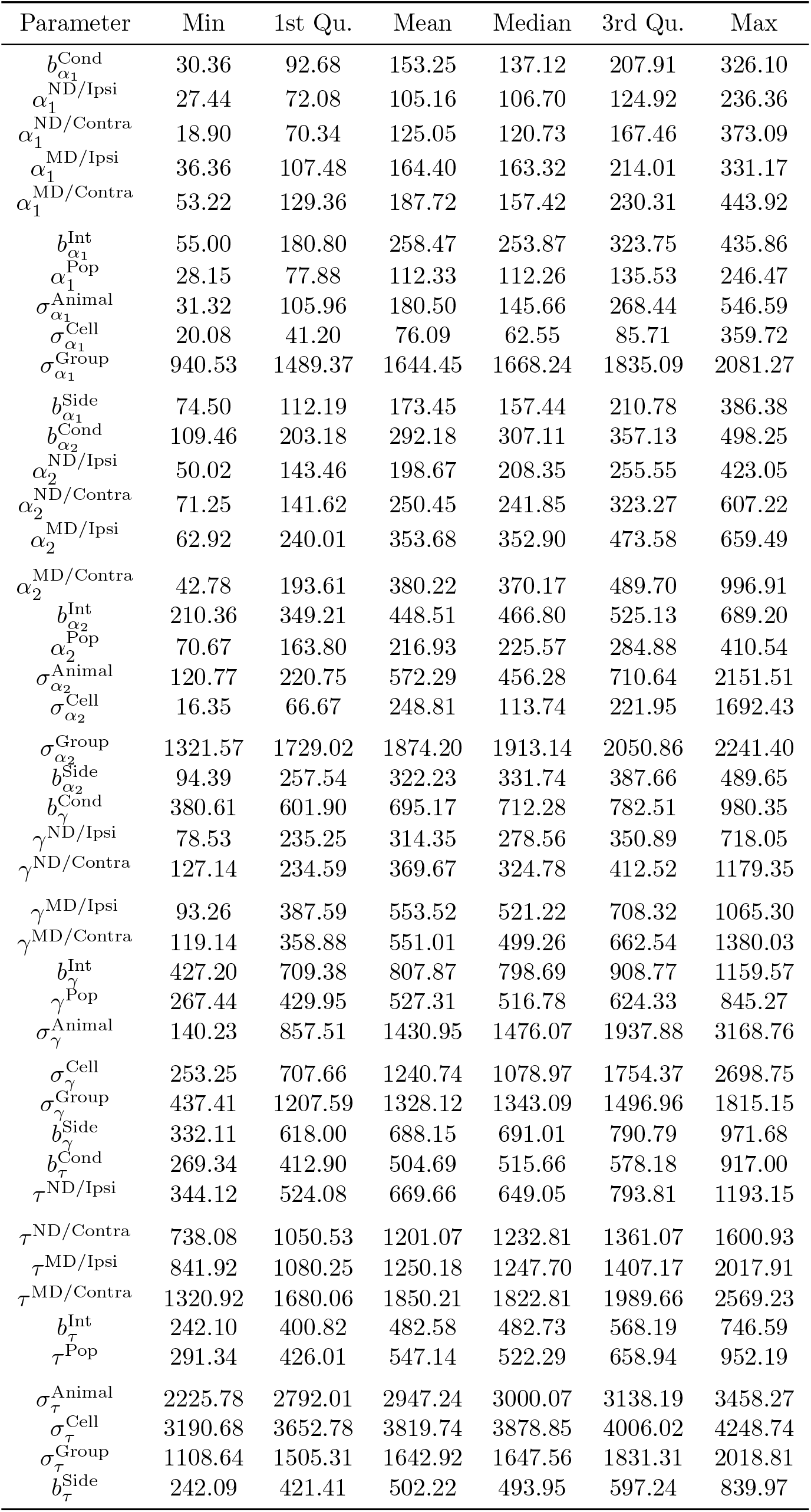
Summary of the effective sample size for parameters at the animal-level in simulation scenario 2. The reported summaries are taken across the 50 simulation runs.

**Table S13.**
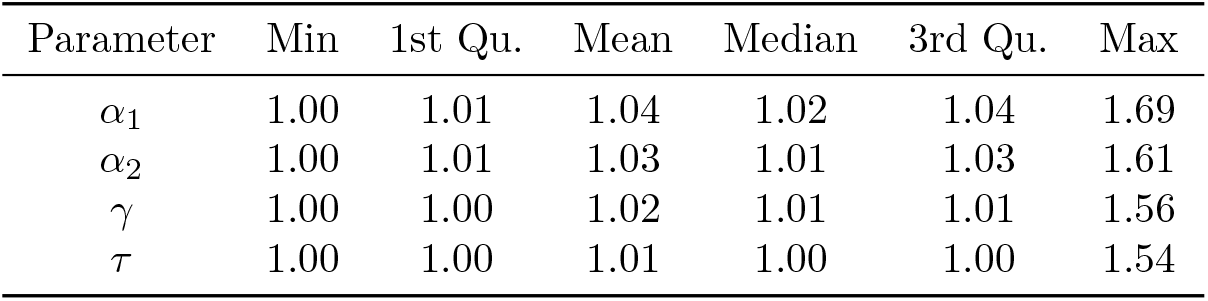
Summary of R-hat at the cell-level for simulation scenario 3. Due to the number of cell level parameters, the reported summaries are taken across the 50 simulation runs AND all cell-level parameters.

**Table S14.** Summary of R-hat at the animal-level for simulation scenario 3. Due to the number of animal level parameters, the reported summaries are taken across the 50 simulation runs AND all animal-level parameters.

| Parameter | Min | 1st Qu. | Mean | Median | 3rd Qu. | Max |
| --- | --- | --- | --- | --- | --- | --- |
| $\alpha_1$ | 1.00 | 1.02 | 1.05 | 1.03 | 1.05 | 1.71 |
| $\alpha_2$ | 1.00 | 1.01 | 1.04 | 1.02 | 1.04 | 1.62 |
| $\gamma$ | 1.00 | 1.01 | 1.03 | 1.02 | 1.03 | 1.59 |
| $\tau$ | 1.00 | 1.00 | 1.01 | 1.00 | 1.00 | 1.55 |

**Table S15.** Summary of R-hat for other model parameters in simulation scenario 3. The reported summaries are taken across the 50 simulation runs.

| Parameter | Min | 1st Qu. | Mean | Median | 3rd Qu. | Max |
| --- | --- | --- | --- | --- | --- | --- |
| $b_{\alpha_1}^{\text{Cond}}$ | 1.01 | 1.02 | 1.03 | 1.03 | 1.04 | 1.10 |
| $\alpha_1^{\text{ND/Ipsi}}$ | 1.02 | 1.03 | 1.06 | 1.04 | 1.06 | 1.62 |
| $\alpha_1^{\text{ND/Contra}}$ | 1.01 | 1.03 | 1.06 | 1.04 | 1.06 | 1.61 |
| $\alpha_1^{\text{MD/Ipsi}}$ | 1.01 | 1.02 | 1.05 | 1.03 | 1.06 | 1.63 |
| $\alpha_1^{\text{MD/Contra}}$ | 1.01 | 1.03 | 1.06 | 1.04 | 1.07 | 1.71 |
| $b_{\alpha_1}^{\text{Int}}$ | 1.00 | 1.01 | 1.02 | 1.02 | 1.03 | 1.10 |
| $\alpha_1^{\text{Pop}}$ | 1.02 | 1.02 | 1.05 | 1.04 | 1.05 | 1.61 |
| $\sigma_{\alpha_1}^{\text{Animal}}$ | 1.00 | 1.02 | 1.04 | 1.03 | 1.06 | 1.19 |
| $\sigma_{\alpha_1}^{\text{Cell}}$ | 1.01 | 1.04 | 1.09 | 1.07 | 1.11 | 1.50 |
| $\sigma_{\alpha_1}^{\text{Group}}$ | 1.00 | 1.00 | 1.00 | 1.00 | 1.00 | 1.01 |
| $b_{\alpha_1}^{\text{Side}}$ | 1.01 | 1.02 | 1.03 | 1.03 | 1.04 | 1.08 |
| $b_{\alpha_2}^{\text{Cond}}$ | 1.00 | 1.01 | 1.02 | 1.01 | 1.02 | 1.08 |
| $\alpha_2^{\text{ND/Ipsi}}$ | 1.00 | 1.02 | 1.04 | 1.03 | 1.04 | 1.55 |
| $\alpha_2^{\text{ND/Contra}}$ | 1.01 | 1.02 | 1.04 | 1.03 | 1.04 | 1.56 |
| $\alpha_2^{\text{MD/Ipsi}}$ | 1.00 | 1.01 | 1.04 | 1.02 | 1.03 | 1.55 |
| $\alpha_2^{\text{MD/Contra}}$ | 1.00 | 1.02 | 1.04 | 1.03 | 1.04 | 1.61 |
| $b_{\alpha_2}^{\text{Int}}$ | 1.00 | 1.01 | 1.02 | 1.01 | 1.02 | 1.15 |
| $\alpha_2^{\text{Pop}}$ | 1.00 | 1.02 | 1.04 | 1.02 | 1.03 | 1.55 |
| $\sigma_{\alpha_2}^{\text{Animal}}$ | 1.00 | 1.01 | 1.03 | 1.02 | 1.03 | 1.24 |
| $\sigma_{\alpha_2}^{\text{Cell}}$ | 1.00 | 1.02 | 1.07 | 1.05 | 1.10 | 1.31 |
| $\sigma_{\alpha_2}^{\text{Group}}$ | 1.00 | 1.00 | 1.00 | 1.00 | 1.00 | 1.01 |
| $b_{\alpha_2}^{\text{Side}}$ | 1.00 | 1.01 | 1.02 | 1.02 | 1.03 | 1.15 |
| $b_{\gamma}^{\text{Cond}}$ | 1.00 | 1.00 | 1.01 | 1.01 | 1.01 | 1.02 |
| $\gamma^{\text{ND/Ipsi}}$ | 1.00 | 1.01 | 1.03 | 1.01 | 1.02 | 1.55 |
| $\gamma^{\text{ND/Contra}}$ | 1.00 | 1.01 | 1.03 | 1.02 | 1.03 | 1.54 |
| $\gamma^{\text{MD/Ipsi}}$ | 1.00 | 1.01 | 1.03 | 1.01 | 1.02 | 1.54 |
| $\gamma^{\text{MD/Contra}}$ | 1.00 | 1.01 | 1.03 | 1.02 | 1.03 | 1.60 |
| $b_{\gamma}^{\text{Int}}$ | 1.00 | 1.00 | 1.01 | 1.01 | 1.01 | 1.02 |
| $\gamma^{\text{Pop}}$ | 1.00 | 1.01 | 1.02 | 1.01 | 1.01 | 1.53 |
| $\sigma_{\gamma}^{\text{Animal}}$ | 1.00 | 1.00 | 1.01 | 1.00 | 1.01 | 1.03 |
| $\sigma_{\gamma}^{\text{Cell}}$ | 1.00 | 1.00 | 1.01 | 1.01 | 1.01 | 1.25 |
| $\sigma_{\gamma}^{\text{Group}}$ | 1.00 | 1.00 | 1.00 | 1.00 | 1.00 | 1.01 |
| $b_{\gamma}^{\text{Side}}$ | 1.00 | 1.00 | 1.01 | 1.01 | 1.01 | 1.02 |
| $b_{\tau}^{\text{Cond}}$ | 1.00 | 1.00 | 1.01 | 1.01 | 1.01 | 1.03 |
| $\tau^{\text{ND/Ipsi}}$ | 1.00 | 1.00 | 1.02 | 1.00 | 1.01 | 1.54 |
| $\tau^{\text{ND/Contra}}$ | 1.00 | 1.00 | 1.01 | 1.00 | 1.00 | 1.53 |
| $\tau^{\text{MD/Ipsi}}$ | 1.00 | 1.00 | 1.01 | 1.00 | 1.00 | 1.53 |
| $\tau^{\text{MD/Contra}}$ | 1.00 | 1.00 | 1.01 | 1.00 | 1.00 | 1.53 |
| $b_{\tau}^{\text{Int}}$ | 1.00 | 1.00 | 1.01 | 1.01 | 1.01 | 1.02 |
| $\tau^{\text{Pop}}$ | 1.00 | 1.00 | 1.02 | 1.01 | 1.01 | 1.53 |
| $\sigma_{\tau}^{\text{Animal}}$ | 1.00 | 1.00 | 1.00 | 1.00 | 1.00 | 1.01 |
| $\sigma_{\tau}^{\text{Cell}}$ | 1.00 | 1.00 | 1.00 | 1.00 | 1.00 | 1.01 |
| $\sigma_{\tau}^{\text{Group}}$ | 1.00 | 1.00 | 1.00 | 1.00 | 1.00 | 1.01 |
| $b_{\tau}^{\text{Side}}$ | 1.00 | 1.00 | 1.01 | 1.01 | 1.01 | 1.02 |

**Table S16.** Summary of the effective sample size for parameters at the cell-level in simulation scenario 3. Due to the number of cell level parameters, the reported summaries are taken across the 50 simulation runs AND all cell-level parameters.

| Parameter | Min | 1st Qu. | Mean | Median | 3rd Qu. | Max |
| --- | --- | --- | --- | --- | --- | --- |
| $\alpha_1$ | 6.37 | 105.14 | 266.43 | 179.50 | 333.79 | 1996.41 |
| $\alpha_2$ | 6.77 | 175.76 | 475.21 | 329.11 | 664.90 | 2544.24 |
| $\gamma$ | 6.98 | 324.76 | 554.77 | 494.92 | 729.60 | 2052.84 |
| $\tau$ | 7.09 | 3051.74 | 3258.37 | 3371.76 | 3634.91 | 4343.75 |

**Table S17.** Summary of the effective sample size for parameters at the animal-level in simulation scenario 3. Due to the number of animal level parameters, the reported summaries are taken across the 50 simulation runs AND all animal-level parameters.

| Parameter | Min | 1st Qu. | Mean | Median | 3rd Qu. | Max |
| --- | --- | --- | --- | --- | --- | --- |
| $\alpha_1$ | 6.31 | 74.13 | 134.93 | 118.40 | 173.72 | 453.28 |
| $\alpha_2$ | 6.77 | 114.65 | 218.24 | 184.71 | 298.38 | 668.99 |
| $\gamma$ | 6.84 | 135.80 | 238.85 | 210.60 | 302.29 | 826.14 |
| $\tau$ | 7.05 | 2132.71 | 2488.94 | 2538.66 | 2931.25 | 4042.59 |

**Table S18.** Summary of the effective sample size for parameters at the animal-level in simulation scenario 3. The reported summaries are taken across the 50 simulation runs.

| Parameter | Min | 1st Qu. | Mean | Median | 3rd Qu. | Max |
| --- | --- | --- | --- | --- | --- | --- |
| $b_{\alpha_1}^{\text{Cond}}$ | 31.04 | 92.73 | 153.92 | 161.76 | 212.65 | 281.61 |
| $\alpha_1^{\text{ND/Ipsi}}$ | 6.73 | 56.52 | 102.19 | 96.29 | 132.79 | 233.45 |
| $\alpha_1^{\text{ND/Contra}}$ | 6.84 | 61.80 | 99.40 | 87.01 | 130.48 | 203.24 |
| $\alpha_1^{\text{MD/Ipsi}}$ | 6.88 | 67.86 | 122.50 | 124.95 | 154.13 | 298.08 |
| $\alpha_1^{\text{MD/Contra}}$ | 6.37 | 52.92 | 95.09 | 83.29 | 125.51 | 246.80 |
| $b_{\alpha_1}^{\text{Int}}$ | 51.05 | 144.16 | 197.91 | 187.39 | 243.57 | 516.20 |
| $\alpha_1^{\text{Pop}}$ | 6.84 | 62.29 | 107.10 | 105.07 | 137.97 | 245.43 |
| $\sigma_{\alpha_1}^{\text{Animal}}$ | 17.44 | 65.91 | 154.00 | 128.62 | 198.65 | 509.31 |
| $\sigma_{\alpha_1}^{\text{Cell}}$ | 7.87 | 32.77 | 60.88 | 50.20 | 82.40 | 166.76 |
| $\sigma_{\alpha_1}^{\text{Group}}$ | 740.27 | 1581.38 | 1694.44 | 1724.93 | 1850.43 | 2063.79 |
| $b_{\alpha_1}^{\text{Side}}$ | 44.42 | 95.87 | 140.75 | 145.11 | 177.61 | 240.60 |
| $b_{\alpha_2}^{\text{Cond}}$ | 45.23 | 191.75 | 265.06 | 251.12 | 346.87 | 532.47 |
| $\alpha_2^{\text{ND/Ipsi}}$ | 7.24 | 115.71 | 166.25 | 159.10 | 215.88 | 357.42 |
| $\alpha_2^{\text{ND/Contra}}$ | 7.12 | 107.58 | 172.50 | 155.55 | 225.90 | 407.53 |
| $\alpha_2^{\text{MD/Ipsi}}$ | 7.07 | 127.17 | 196.45 | 174.31 | 273.59 | 444.71 |
| $\alpha_2^{\text{MD/Contra}}$ | 6.88 | 105.23 | 160.45 | 142.54 | 205.65 | 368.04 |
| $b_{\alpha_2}^{\text{Int}}$ | 19.03 | 231.75 | 317.62 | 327.18 | 383.75 | 536.61 |
| $\alpha_2^{\text{Pop}}$ | 7.21 | 132.62 | 183.76 | 181.75 | 245.69 | 370.27 |
| $\sigma_{\alpha_2}^{\text{Animal}}$ | 12.89 | 138.24 | 314.98 | 268.23 | 460.51 | 998.17 |
| $\sigma_{\alpha_2}^{\text{Cell}}$ | 11.13 | 45.17 | 141.55 | 78.23 | 179.03 | 714.92 |
| $\sigma_{\alpha_2}^{\text{Group}}$ | 1339.78 | 1738.93 | 1856.44 | 1865.71 | 2024.53 | 2227.03 |
| $b_{\alpha_2}^{\text{Side}}$ | 18.63 | 180.41 | 245.35 | 254.30 | 306.26 | 406.24 |
| $b_{\gamma}^{\text{Cond}}$ | 299.72 | 552.90 | 669.84 | 637.19 | 793.57 | 1135.51 |
| $\gamma^{\text{ND/Ipsi}}$ | 7.14 | 170.36 | 305.08 | 295.77 | 424.26 | 716.68 |
| $\gamma^{\text{ND/Contra}}$ | 7.15 | 140.78 | 257.54 | 231.93 | 341.84 | 768.40 |
| $\gamma^{\text{MD/Ipsi}}$ | 7.21 | 206.00 | 327.74 | 289.32 | 424.09 | 773.41 |
| $\gamma^{\text{MD/Contra}}$ | 6.82 | 157.19 | 229.59 | 204.39 | 289.71 | 620.36 |
| $b_{\gamma}^{\text{Int}}$ | 415.71 | 612.89 | 705.10 | 716.91 | 816.42 | 1068.36 |
| $\gamma^{\text{Pop}}$ | 7.23 | 406.73 | 510.00 | 507.27 | 612.18 | 992.78 |
| $\sigma_{\gamma}^{\text{Animal}}$ | 91.82 | 423.66 | 1076.68 | 857.35 | 1495.07 | 3066.69 |
| $\sigma_{\gamma}^{\text{Cell}}$ | 11.74 | 558.06 | 880.76 | 787.26 | 1076.20 | 3325.31 |
| $\sigma_{\gamma}^{\text{Group}}$ | 841.15 | 1199.09 | 1348.37 | 1350.29 | 1492.69 | 1692.14 |
| $b_{\gamma}^{\text{Side}}$ | 251.02 | 488.17 | 593.15 | 591.14 | 687.93 | 978.95 |
| $b_{\tau}^{\text{Cond}}$ | 262.67 | 415.95 | 523.44 | 517.90 | 617.07 | 814.72 |
| $\tau^{\text{ND/Ipsi}}$ | 7.19 | 545.86 | 672.43 | 654.53 | 812.59 | 1054.68 |
| $\tau^{\text{ND/Contra}}$ | 7.25 | 1018.42 | 1175.38 | 1170.70 | 1322.25 | 1850.04 |
| $\tau^{\text{MD/Ipsi}}$ | 7.22 | 1052.61 | 1216.44 | 1280.48 | 1419.59 | 1729.18 |
| $\tau^{\text{MD/Contra}}$ | 7.23 | 1526.34 | 1630.46 | 1638.73 | 1811.07 | 2146.61 |
| $b_{\tau}^{\text{Int}}$ | 289.40 | 422.75 | 501.08 | 492.16 | 590.48 | 706.62 |
| $\tau^{\text{Pop}}$ | 7.38 | 452.55 | 551.84 | 515.18 | 668.62 | 818.51 |
| $\sigma_{\tau}^{\text{Animal}}$ | 2259.42 | 2663.99 | 2845.03 | 2873.41 | 3007.70 | 3561.13 |
| $\sigma_{\tau}^{\text{Cell}}$ | 2926.08 | 3426.54 | 3678.84 | 3791.55 | 3948.19 | 4306.71 |
| $\sigma_{\tau}^{\text{Group}}$ | 966.21 | 1505.71 | 1646.89 | 1648.96 | 1775.15 | 2036.35 |
| $b_{\tau}^{\text{Side}}$ | 313.18 | 430.21 | 527.54 | 507.12 | 607.66 | 809.99 |

**Table S19.**
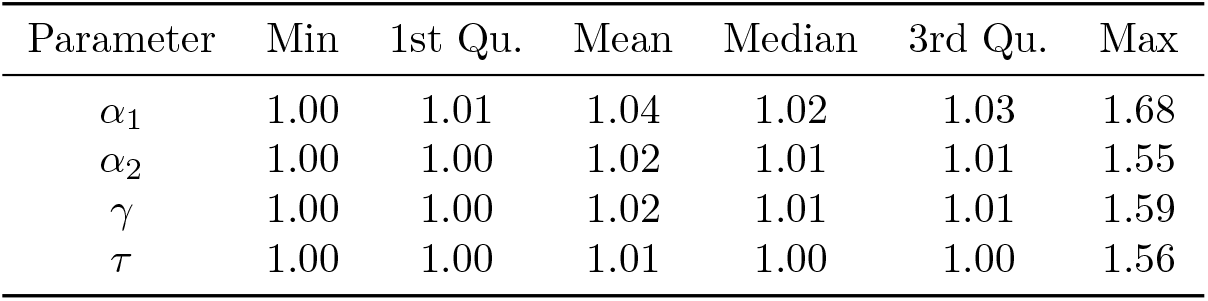
Summary of R-hat at the cell-level for simulation scenario 4. Due to the number of cell level parameters, the reported summaries are taken across the 50 simulation runs AND all cell-level parameters.

| Parameter | Min | 1st Qu. | Mean | Median | 3rd Qu. | Max |
| --- | --- | --- | --- | --- | --- | --- |
| $\alpha_1$ | 1.00 | 1.01 | 1.04 | 1.02 | 1.03 | 1.68 |
| $\alpha_2$ | 1.00 | 1.00 | 1.02 | 1.01 | 1.01 | 1.55 |
| $\gamma$ | 1.00 | 1.00 | 1.02 | 1.01 | 1.01 | 1.59 |
| $\tau$ | 1.00 | 1.00 | 1.01 | 1.00 | 1.00 | 1.56 |

**Table S20.** Summary of R-hat at the animal-level for simulation scenario 4. Due to the number of animal level parameters, the reported summaries are taken across the 50 simulation runs AND all animal-level parameters.

| Parameter | Min | 1st Qu. | Mean | Median | 3rd Qu. | Max |
| --- | --- | --- | --- | --- | --- | --- |
| $\alpha_1$ | 1.00 | 1.02 | 1.06 | 1.03 | 1.05 | 1.69 |
| $\alpha_2$ | 1.00 | 1.01 | 1.03 | 1.01 | 1.02 | 1.58 |
| $\gamma$ | 1.00 | 1.01 | 1.03 | 1.01 | 1.02 | 1.61 |
| $\tau$ | 1.00 | 1.00 | 1.01 | 1.00 | 1.00 | 1.57 |

**Table S21.** Summary of R-hat for other model parameters in simulation scenario 4. The reported summaries are taken across the 50 simulation runs.

| Parameter | Min | 1st Qu. | Mean | Median | 3rd Qu. | Max |
| --- | --- | --- | --- | --- | --- | --- |
| $b_{\alpha_1}^{\text{Cond}}$ | 1.00 | 1.02 | 1.04 | 1.03 | 1.04 | 1.22 |
| $\alpha_1^{\text{ND/Ipsi}}$ | 1.01 | 1.03 | 1.07 | 1.04 | 1.07 | 1.63 |
| $\alpha_1^{\text{ND/Contra}}$ | 1.01 | 1.03 | 1.07 | 1.05 | 1.06 | 1.69 |
| $\alpha_1^{\text{MD/Ipsi}}$ | 1.00 | 1.02 | 1.06 | 1.04 | 1.04 | 1.62 |
| $\alpha_1^{\text{MD/Contra}}$ | 1.00 | 1.01 | 1.05 | 1.02 | 1.04 | 1.62 |
| $b_{\alpha_1}^{\text{Int}}$ | 1.00 | 1.01 | 1.03 | 1.02 | 1.03 | 1.14 |
| $\alpha_1^{\text{Pop}}$ | 1.01 | 1.03 | 1.07 | 1.04 | 1.06 | 1.63 |
| $\sigma_{\alpha_1}^{\text{Animal}}$ | 1.00 | 1.01 | 1.04 | 1.03 | 1.05 | 1.35 |
| $\sigma_{\alpha_1}^{\text{Cell}}$ | 1.01 | 1.04 | 1.09 | 1.06 | 1.11 | 1.61 |
| $\sigma_{\alpha_1}^{\text{Group}}$ | 1.00 | 1.00 | 1.00 | 1.00 | 1.00 | 1.01 |
| $b_{\alpha_1}^{\text{Side}}$ | 1.01 | 1.02 | 1.04 | 1.03 | 1.04 | 1.20 |
| $b_{\alpha_2}^{\text{Cond}}$ | 1.00 | 1.01 | 1.02 | 1.01 | 1.02 | 1.08 |
| $\alpha_2^{\text{ND/Ipsi}}$ | 1.00 | 1.01 | 1.04 | 1.02 | 1.03 | 1.55 |
| $\alpha_2^{\text{ND/Contra}}$ | 1.01 | 1.01 | 1.03 | 1.02 | 1.03 | 1.59 |
| $\alpha_2^{\text{MD/Ipsi}}$ | 1.00 | 1.01 | 1.03 | 1.01 | 1.02 | 1.58 |
| $\alpha_2^{\text{MD/Contra}}$ | 1.00 | 1.01 | 1.02 | 1.01 | 1.02 | 1.58 |
| $b_{\alpha_2}^{\text{Int}}$ | 1.00 | 1.01 | 1.01 | 1.01 | 1.01 | 1.03 |
| $\alpha_2^{\text{Pop}}$ | 1.01 | 1.01 | 1.03 | 1.02 | 1.03 | 1.55 |
| $\sigma_{\alpha_2}^{\text{Animal}}$ | 1.00 | 1.00 | 1.01 | 1.01 | 1.01 | 1.07 |
| $\sigma_{\alpha_2}^{\text{Cell}}$ | 1.00 | 1.01 | 1.05 | 1.03 | 1.04 | 1.42 |
| $\sigma_{\alpha_2}^{\text{Group}}$ | 1.00 | 1.00 | 1.00 | 1.00 | 1.00 | 1.01 |
| $b_{\alpha_2}^{\text{Side}}$ | 1.00 | 1.01 | 1.02 | 1.02 | 1.02 | 1.08 |
| $b_{\gamma}^{\text{Cond}}$ | 1.00 | 1.00 | 1.01 | 1.01 | 1.01 | 1.02 |
| $\gamma^{\text{ND/Ipsi}}$ | 1.00 | 1.01 | 1.03 | 1.01 | 1.02 | 1.55 |
| $\gamma^{\text{ND/Contra}}$ | 1.01 | 1.01 | 1.03 | 1.02 | 1.02 | 1.56 |
| $\gamma^{\text{MD/Ipsi}}$ | 1.00 | 1.01 | 1.02 | 1.01 | 1.02 | 1.57 |
| $\gamma^{\text{MD/Contra}}$ | 1.00 | 1.00 | 1.02 | 1.01 | 1.01 | 1.57 |
| $b_{\gamma}^{\text{Int}}$ | 1.00 | 1.00 | 1.01 | 1.00 | 1.01 | 1.02 |
| $\gamma^{\text{Pop}}$ | 1.00 | 1.00 | 1.02 | 1.01 | 1.01 | 1.45 |
| $\sigma_{\gamma}^{\text{Animal}}$ | 1.00 | 1.00 | 1.00 | 1.00 | 1.01 | 1.05 |
| $\sigma_{\gamma}^{\text{Cell}}$ | 1.00 | 1.00 | 1.01 | 1.00 | 1.00 | 1.15 |
| $\sigma_{\gamma}^{\text{Group}}$ | 1.00 | 1.00 | 1.00 | 1.00 | 1.00 | 1.01 |
| $b_{\gamma}^{\text{Side}}$ | 1.00 | 1.00 | 1.01 | 1.01 | 1.01 | 1.02 |
| $b_{\tau}^{\text{Cond}}$ | 1.00 | 1.00 | 1.01 | 1.01 | 1.01 | 1.06 |
| $\tau^{\text{ND/Ipsi}}$ | 1.00 | 1.00 | 1.02 | 1.00 | 1.01 | 1.54 |
| $\tau^{\text{ND/Contra}}$ | 1.00 | 1.00 | 1.01 | 1.00 | 1.00 | 1.54 |
| $\tau^{\text{MD/Ipsi}}$ | 1.00 | 1.00 | 1.01 | 1.00 | 1.00 | 1.53 |
| $\tau^{\text{MD/Contra}}$ | 1.00 | 1.00 | 1.01 | 1.00 | 1.00 | 1.53 |
| $b_{\tau}^{\text{Int}}$ | 1.00 | 1.00 | 1.01 | 1.01 | 1.01 | 1.02 |
| $\tau^{\text{Pop}}$ | 1.00 | 1.00 | 1.02 | 1.01 | 1.01 | 1.53 |
| $\sigma_{\tau}^{\text{Animal}}$ | 1.00 | 1.00 | 1.00 | 1.00 | 1.00 | 1.01 |
| $\sigma_{\tau}^{\text{Cell}}$ | 1.00 | 1.00 | 1.00 | 1.00 | 1.00 | 1.02 |
| $\sigma_{\tau}^{\text{Group}}$ | 1.00 | 1.00 | 1.00 | 1.00 | 1.00 | 1.01 |
| $b_{\tau}^{\text{Side}}$ | 1.00 | 1.00 | 1.01 | 1.01 | 1.01 | 1.02 |

**Table S22.** Summary of the effective sample size for parameters at the cell-level in simulation scenario 4. Due to the number of cell level parameters, the reported summaries are taken across the 50 simulation runs AND all cell-level parameters.

| Parameter | Min | 1st Qu. | Mean | Median | 3rd Qu. | Max |
| --- | --- | --- | --- | --- | --- | --- |
| $\alpha_1$ | 6.39 | 129.23 | 298.57 | 220.21 | 382.89 | 1920.85 |
| $\alpha_2$ | 6.99 | 328.06 | 691.34 | 594.05 | 930.17 | 3319.50 |
| $\gamma$ | 6.78 | 504.60 | 754.90 | 716.64 | 963.59 | 2528.36 |
| $\tau$ | 6.93 | 3224.80 | 3359.88 | 3506.84 | 3730.98 | 4269.48 |

**Table S23.**
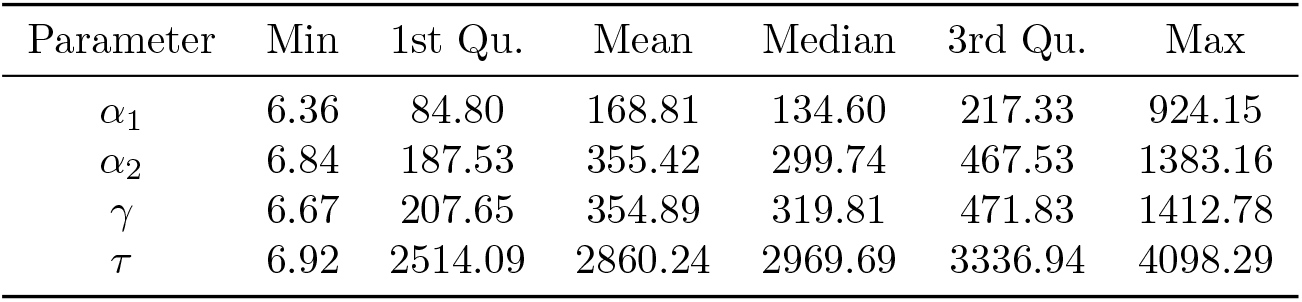
Summary of the effective sample size for parameters at the animal-level in simulation scenario 4. Due to the number of animal level parameters, the reported summaries are taken across the 50 simulation runs AND all animal-level parameters.

**Table S24.**
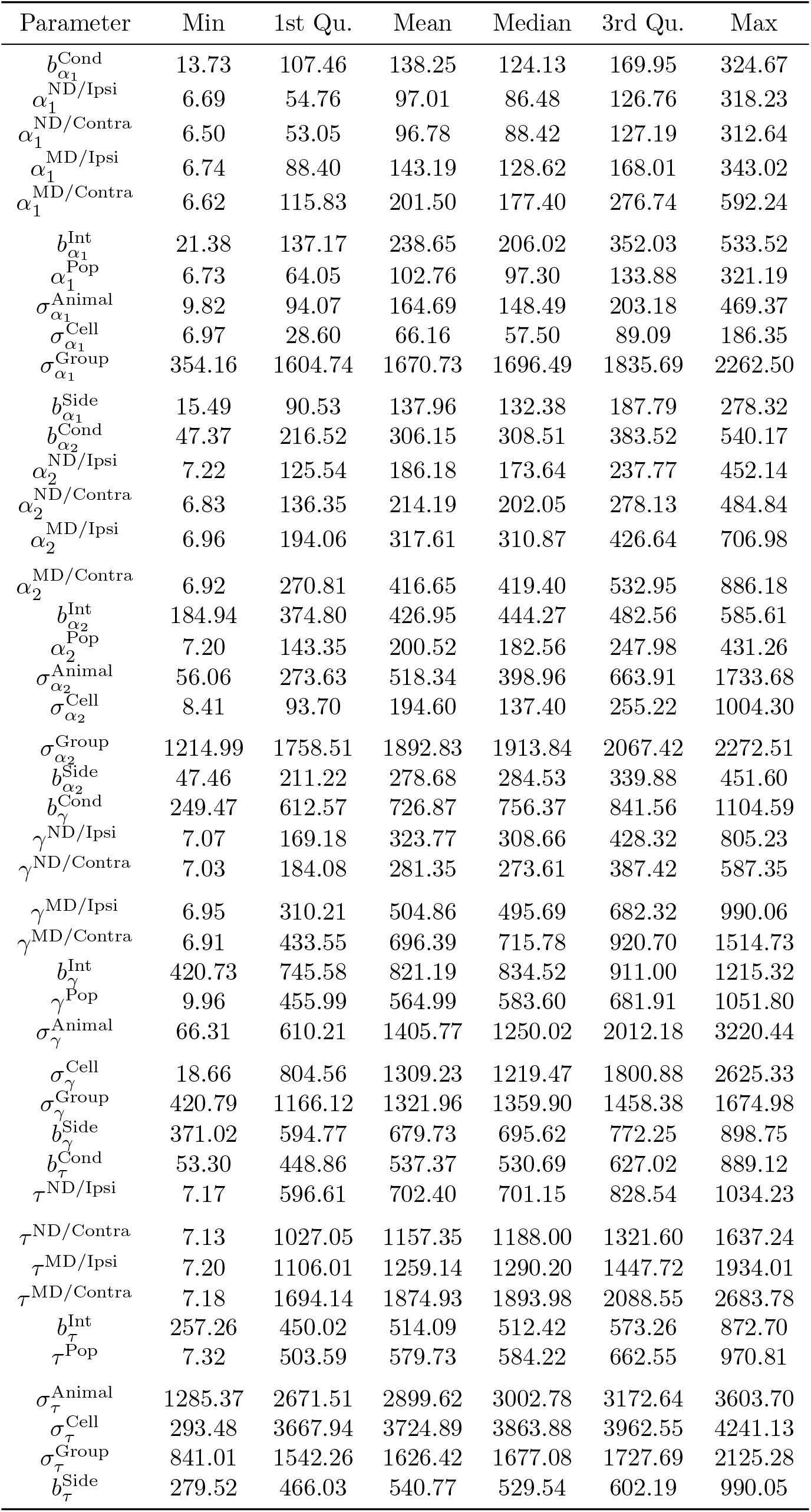
Summary of the effective sample size for parameters at the animal-level in simulation scenario 4. The reported summaries are taken across the 50 simulation runs.

**Table S25.** Summary of R-hat at the cell-level for simulation scenario 5. Due to the number of cell level parameters, the reported summaries are taken across the 50 simulation runs AND all cell-level parameters.

| Parameter | Min | 1st Qu. | Mean | Median | 3rd Qu. | Max |
| --- | --- | --- | --- | --- | --- | --- |
| $\alpha_1$ | 1.00 | 1.01 | 1.03 | 1.02 | 1.03 | 1.27 |
| $\alpha_2$ | 1.00 | 1.00 | 1.01 | 1.00 | 1.01 | 1.14 |
| $\gamma$ | 1.00 | 1.00 | 1.01 | 1.00 | 1.01 | 1.06 |
| $\tau$ | 1.00 | 1.00 | 1.00 | 1.00 | 1.00 | 1.01 |

**Table S26.**
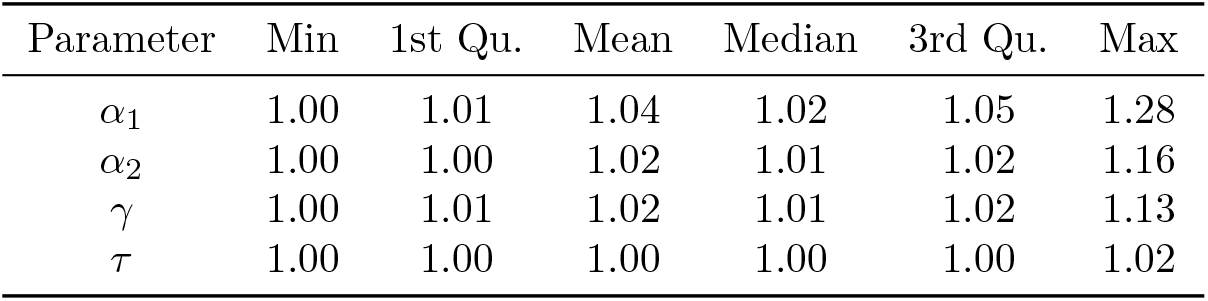
Summary of R-hat at the animal-level for simulation scenario 5. Due to the number of animal level parameters, the reported summaries are taken across the 50 simulation runs AND all animal-level parameters.

**Table S27.** Summary of R-hat for other model parameters in simulation scenario 5. The reported summaries are taken across the 50 simulation runs.

| Parameter | Min | 1st Qu. | Mean | Median | 3rd Qu. | Max |
| --- | --- | --- | --- | --- | --- | --- |
| $b_{\alpha_1}^{\text{Cond}}$ | 1.01 | 1.01 | 1.03 | 1.02 | 1.04 | 1.13 |
| $\alpha_1^{\text{ND/Ipsi}}$ | 1.01 | 1.02 | 1.04 | 1.04 | 1.06 | 1.22 |
| $\alpha_1^{\text{ND/Contra}}$ | 1.00 | 1.02 | 1.03 | 1.03 | 1.05 | 1.09 |
| $\alpha_1^{\text{MD/Ipsi}}$ | 1.00 | 1.01 | 1.02 | 1.02 | 1.04 | 1.08 |
| $\alpha_1^{\text{MD/Contra}}$ | 1.00 | 1.01 | 1.02 | 1.01 | 1.03 | 1.15 |
| $b_{\alpha_1}^{\text{Int}}$ | 1.00 | 1.01 | 1.02 | 1.01 | 1.02 | 1.07 |
| $\alpha_1^{\text{Pop}}$ | 1.01 | 1.02 | 1.04 | 1.03 | 1.05 | 1.17 |
| $\sigma_{\alpha_1}^{\text{Animal}}$ | 1.00 | 1.01 | 1.04 | 1.03 | 1.06 | 1.11 |
| $\sigma_{\alpha_1}^{\text{Cell}}$ | 1.01 | 1.04 | 1.11 | 1.07 | 1.14 | 1.49 |
| $\sigma_{\alpha_1}^{\text{Group}}$ | 1.00 | 1.00 | 1.00 | 1.00 | 1.00 | 1.00 |
| $b_{\alpha_1}^{\text{Side}}$ | 1.00 | 1.01 | 1.03 | 1.02 | 1.03 | 1.09 |
| $b_{\alpha_2}^{\text{Cond}}$ | 1.00 | 1.01 | 1.01 | 1.01 | 1.02 | 1.03 |
| $\alpha_2^{\text{ND/Ipsi}}$ | 1.00 | 1.01 | 1.02 | 1.02 | 1.02 | 1.05 |
| $\alpha_2^{\text{ND/Contra}}$ | 1.00 | 1.01 | 1.02 | 1.01 | 1.02 | 1.06 |
| $\alpha_2^{\text{MD/Ipsi}}$ | 1.00 | 1.00 | 1.01 | 1.01 | 1.01 | 1.06 |
| $\alpha_2^{\text{MD/Contra}}$ | 1.00 | 1.00 | 1.01 | 1.01 | 1.01 | 1.05 |
| $b_{\alpha_2}^{\text{Int}}$ | 1.00 | 1.00 | 1.01 | 1.01 | 1.01 | 1.04 |
| $\alpha_2^{\text{Pop}}$ | 1.00 | 1.01 | 1.02 | 1.01 | 1.02 | 1.05 |
| $\sigma_{\alpha_2}^{\text{Animal}}$ | 1.00 | 1.00 | 1.01 | 1.01 | 1.02 | 1.09 |
| $\sigma_{\alpha_2}^{\text{Cell}}$ | 1.00 | 1.01 | 1.04 | 1.02 | 1.04 | 1.35 |
| $\sigma_{\alpha_2}^{\text{Group}}$ | 1.00 | 1.00 | 1.00 | 1.00 | 1.00 | 1.01 |
| $b_{\alpha_2}^{\text{Side}}$ | 1.00 | 1.01 | 1.01 | 1.01 | 1.01 | 1.06 |
| $b_{\gamma}^{\text{Cond}}$ | 1.00 | 1.00 | 1.01 | 1.00 | 1.01 | 1.02 |
| $\gamma^{\text{ND/Ipsi}}$ | 1.00 | 1.01 | 1.01 | 1.01 | 1.01 | 1.07 |
| $\gamma^{\text{ND/Contra}}$ | 1.00 | 1.01 | 1.01 | 1.01 | 1.01 | 1.04 |
| $\gamma^{\text{MD/Ipsi}}$ | 1.00 | 1.00 | 1.01 | 1.00 | 1.01 | 1.05 |
| $\gamma^{\text{MD/Contra}}$ | 1.00 | 1.00 | 1.01 | 1.00 | 1.01 | 1.02 |
| $b_{\gamma}^{\text{Int}}$ | 1.00 | 1.00 | 1.00 | 1.00 | 1.01 | 1.01 |
| $\gamma^{\text{Pop}}$ | 1.00 | 1.00 | 1.01 | 1.01 | 1.01 | 1.02 |
| $\sigma_{\gamma}^{\text{Animal}}$ | 1.00 | 1.00 | 1.00 | 1.00 | 1.00 | 1.05 |
| $\sigma_{\gamma}^{\text{Cell}}$ | 1.00 | 1.00 | 1.01 | 1.00 | 1.01 | 1.03 |
| $\sigma_{\gamma}^{\text{Group}}$ | 1.00 | 1.00 | 1.00 | 1.00 | 1.00 | 1.01 |
| $b_{\gamma}^{\text{Side}}$ | 1.00 | 1.00 | 1.01 | 1.01 | 1.01 | 1.02 |
| $b_{\tau}^{\text{Cond}}$ | 1.00 | 1.00 | 1.01 | 1.01 | 1.01 | 1.02 |
| $\tau^{\text{ND/Ipsi}}$ | 1.00 | 1.00 | 1.01 | 1.01 | 1.01 | 1.01 |
| $\tau^{\text{ND/Contra}}$ | 1.00 | 1.00 | 1.00 | 1.00 | 1.00 | 1.01 |
| $\tau^{\text{MD/Ipsi}}$ | 1.00 | 1.00 | 1.00 | 1.00 | 1.00 | 1.01 |
| $\tau^{\text{MD/Contra}}$ | 1.00 | 1.00 | 1.00 | 1.00 | 1.00 | 1.01 |
| $b_{\tau}^{\text{Int}}$ | 1.00 | 1.01 | 1.01 | 1.01 | 1.01 | 1.02 |
| $\tau^{\text{Pop}}$ | 1.00 | 1.00 | 1.01 | 1.01 | 1.01 | 1.02 |
| $\sigma_{\tau}^{\text{Animal}}$ | 1.00 | 1.00 | 1.00 | 1.00 | 1.00 | 1.00 |
| $\sigma_{\tau}^{\text{Cell}}$ | 1.00 | 1.00 | 1.00 | 1.00 | 1.00 | 1.00 |
| $\sigma_{\tau}^{\text{Group}}$ | 1.00 | 1.00 | 1.00 | 1.00 | 1.00 | 1.01 |
| $b_{\tau}^{\text{Side}}$ | 1.00 | 1.00 | 1.01 | 1.01 | 1.01 | 1.02 |

**Table S28.** Summary of the effective sample size for parameters at the cell-level in simulation scenario 5. Due to the number of cell level parameters, the reported summaries are taken across the 50 simulation runs AND all cell-level parameters.

| Parameter | Min | 1st Qu. | Mean | Median | 3rd Qu. | Max |
| --- | --- | --- | --- | --- | --- | --- |
| $\alpha_1$ | 11.11 | 141.74 | 486.34 | 288.16 | 700.77 | 3040.43 |
| $\alpha_2$ | 19.97 | 599.73 | 1189.70 | 1142.90 | 1671.86 | 3681.30 |
| $\gamma$ | 45.02 | 789.45 | 1226.08 | 1181.48 | 1612.85 | 3428.68 |
| $\tau$ | 1567.24 | 3514.99 | 3672.27 | 3727.87 | 3880.35 | 4455.70 |

**Table S29.** Summary of the effective sample size for parameters at the animal-level in simulation scenario 5. Due to the number of animal level parameters, the reported summaries are taken across the 50 simulation runs AND all animal-level parameters.

| Parameter | Min | 1st Qu. | Mean | Median | 3rd Qu. | Max |
| --- | --- | --- | --- | --- | --- | --- |
| $\alpha_1$ | 11.06 | 86.95 | 236.03 | 161.30 | 317.53 | 1155.84 |
| $\alpha_2$ | 17.95 | 254.59 | 495.23 | 429.68 | 658.39 | 1840.49 |
| $\gamma$ | 20.70 | 213.24 | 426.90 | 356.55 | 578.78 | 1567.86 |
| $\tau$ | 350.33 | 2526.58 | 2937.82 | 3059.34 | 3418.69 | 4166.47 |

**Table S30.**
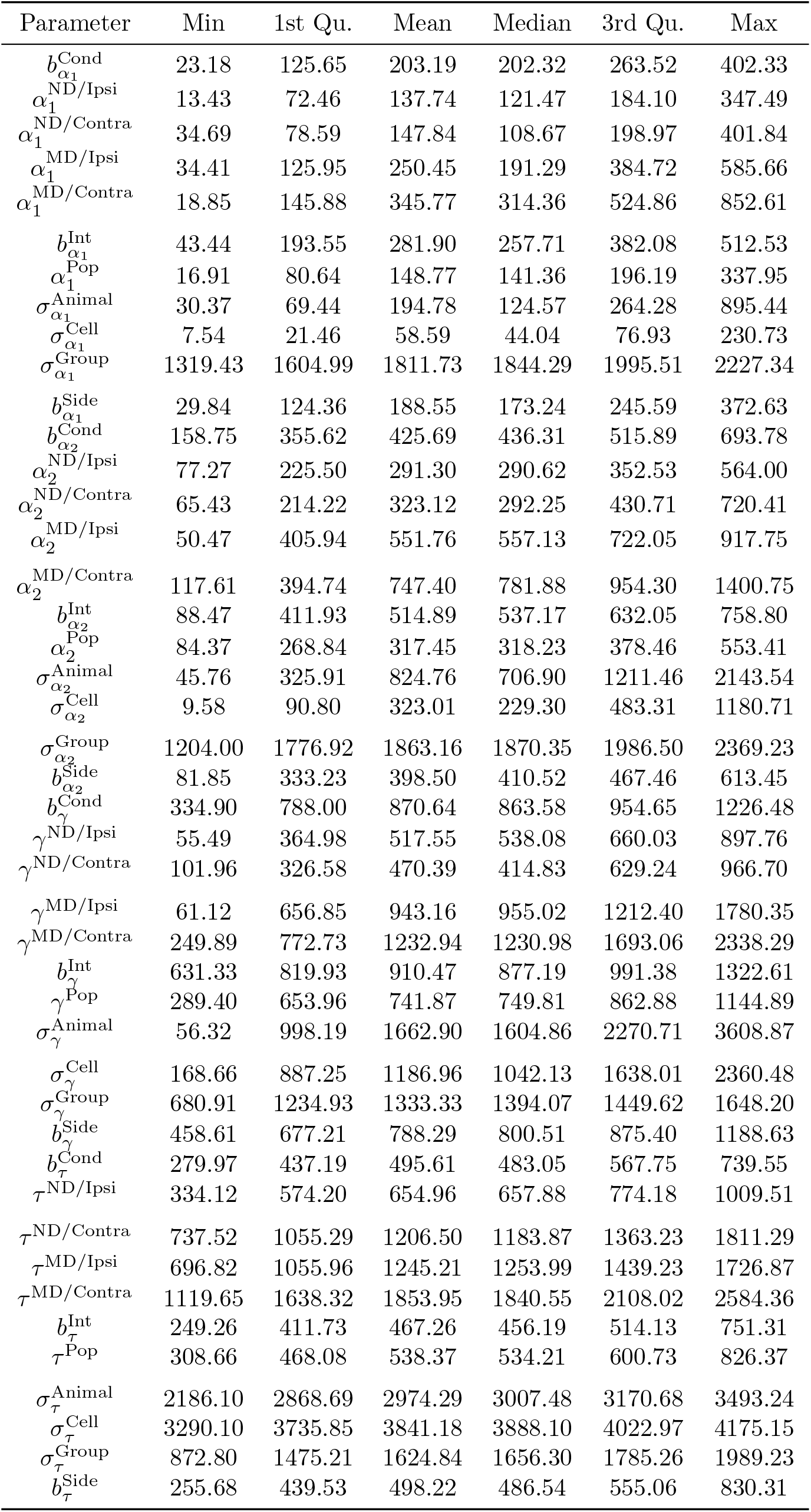
Summary of the effective sample size for parameters at the animal-level in simulation scenario 5. The reported summaries are taken across the 50 simulation runs.

**Table S31.** Summary of R-hat at the cell-level for simulation scenario 6. Due to the number of cell level parameters, the reported summaries are taken across the 50 simulation runs AND all cell-level parameters.

| Parameter | Min | 1st Qu. | Mean | Median | 3rd Qu. | Max |
| --- | --- | --- | --- | --- | --- | --- |
| $\alpha_1$ | 1.00 | 1.01 | 1.04 | 1.02 | 1.03 | 1.69 |
| $\alpha_2$ | 1.00 | 1.00 | 1.02 | 1.01 | 1.01 | 1.59 |
| $\gamma$ | 1.00 | 1.00 | 1.02 | 1.01 | 1.01 | 1.56 |
| $\tau$ | 1.00 | 1.00 | 1.01 | 1.00 | 1.00 | 1.53 |

**Table S32.** Summary of R-hat at the animal-level for simulation scenario 6. Due to the number of animal level parameters, the reported summaries are taken across the 50 simulation runs AND all animal-level parameters.

| Parameter | Min | 1st Qu. | Mean | Median | 3rd Qu. | Max |
| --- | --- | --- | --- | --- | --- | --- |
| $\alpha_1$ | 1.00 | 1.01 | 1.05 | 1.03 | 1.04 | 1.70 |
| $\alpha_2$ | 1.00 | 1.01 | 1.03 | 1.01 | 1.02 | 1.61 |
| $\gamma$ | 1.00 | 1.01 | 1.03 | 1.01 | 1.02 | 1.60 |
| $\tau$ | 1.00 | 1.00 | 1.03 | 1.00 | 1.00 | 1.59 |

**Table S33.**
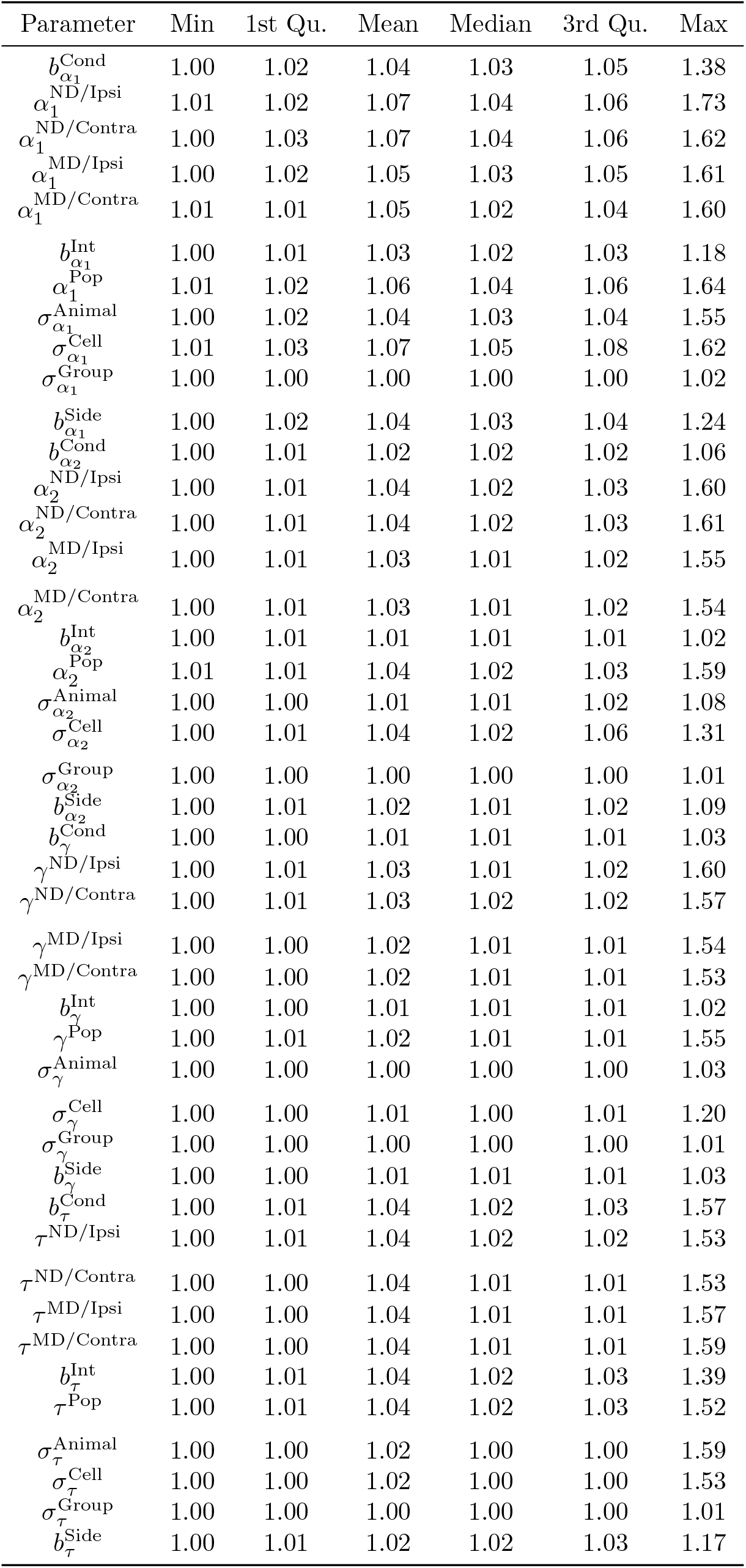
Summary of R-hat for other model parameters in simulation scenario 6. The reported summaries are taken across the 50 simulation runs.

**Table S34.**
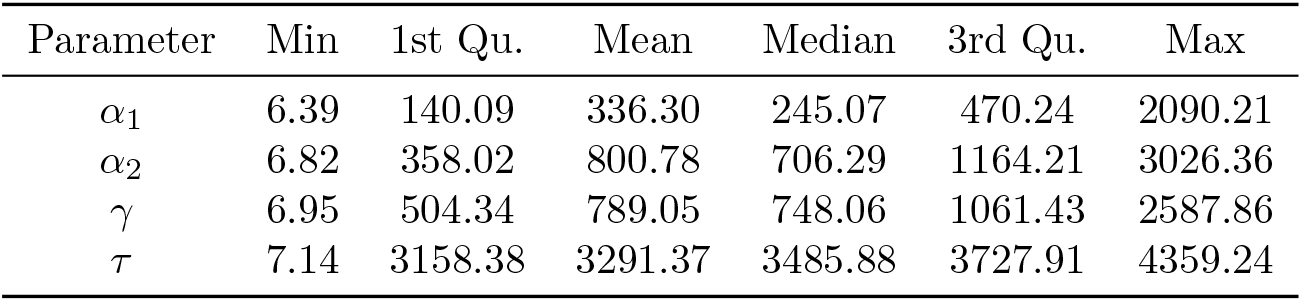
Summary of the effective sample size for parameters at the cell-level in simulation scenario 6. Due to the number of cell level parameters, the reported summaries are taken across the 50 simulation runs AND all cell-level parameters.

| Parameter | Min | 1st Qu. | Mean | Median | 3rd Qu. | Max |
| --- | --- | --- | --- | --- | --- | --- |
| $\alpha_1$ | 6.39 | 140.09 | 336.30 | 245.07 | 470.24 | 2090.21 |
| $\alpha_2$ | 6.82 | 358.02 | 800.78 | 706.29 | 1164.21 | 3026.36 |
| $\gamma$ | 6.95 | 504.34 | 789.05 | 748.06 | 1061.43 | 2587.86 |
| $\tau$ | 7.14 | 3158.38 | 3291.37 | 3485.88 | 3727.91 | 4359.24 |

**Table S35.** Summary of the effective sample size for parameters at the animal-level in simulation scenario 6. Due to the number of animal level parameters, the reported summaries are taken across the 50 simulation runs AND all animal-level parameters.

| Parameter | Min | 1st Qu. | Mean | Median | 3rd Qu. | Max |
| --- | --- | --- | --- | --- | --- | --- |
| $\alpha_1$ | 6.35 | 93.95 | 178.60 | 145.10 | 237.75 | 882.78 |
| $\alpha_2$ | 6.71 | 187.11 | 358.75 | 317.39 | 489.30 | 1397.01 |
| $\gamma$ | 6.79 | 199.43 | 370.92 | 335.35 | 518.86 | 1148.44 |
| $\tau$ | 6.77 | 2579.49 | 2850.93 | 3072.70 | 3454.64 | 4136.71 |

**Table S36.**
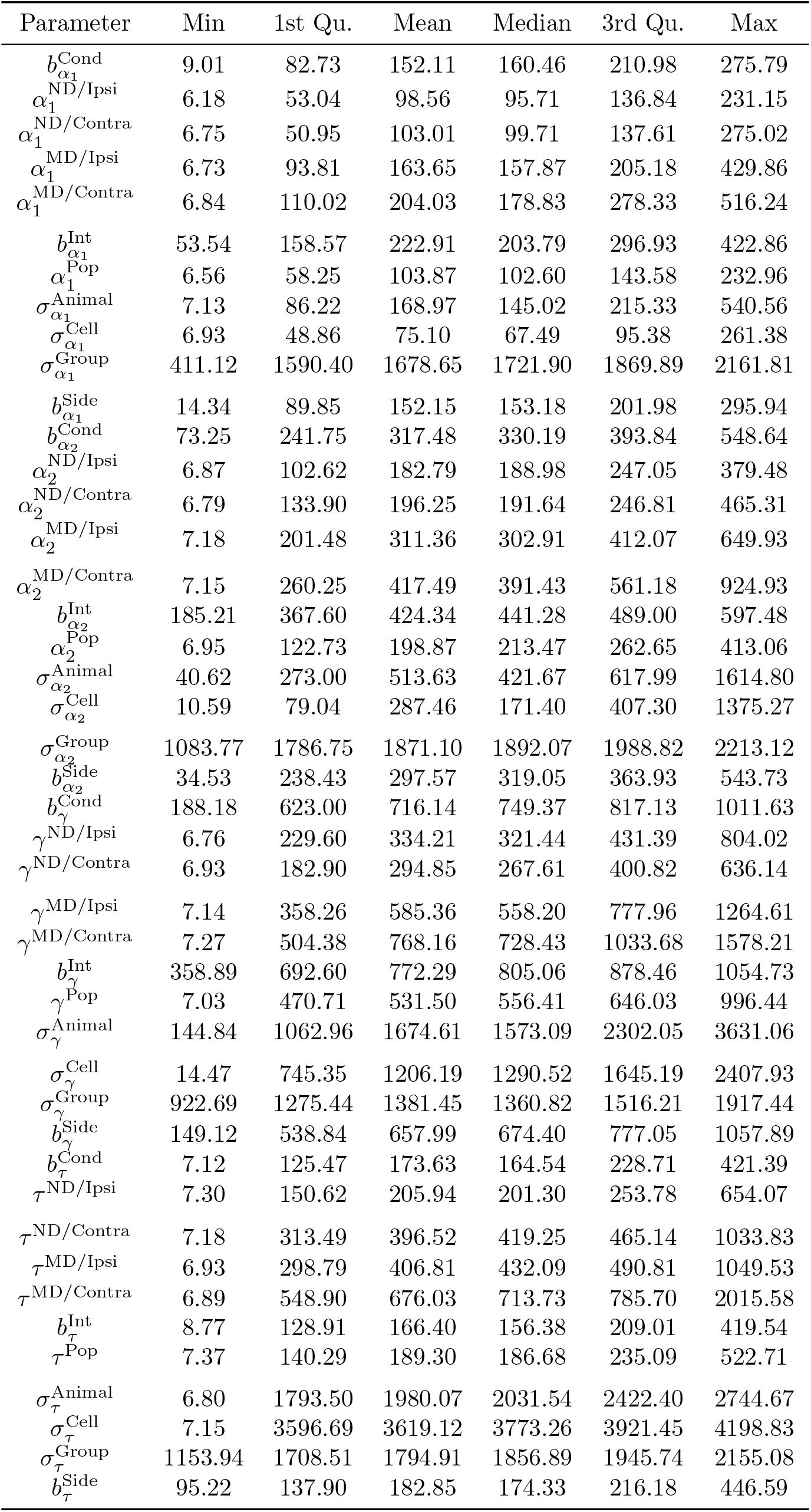
Summary of the effective sample size for parameters at the animal-level in simulation scenario 6. The reported summaries are taken across the 50 simulation runs.

## Notes

https://github.com/vonkaenelerik/hierarchical_sholl

### Competing Interest Statement

The authors have declared no competing interest.

